# High-throughput tandem-microwell assay for ammonia repositions FDA-Approved drugs to *Helicobacter pylori* infection

**DOI:** 10.1101/2021.01.05.425432

**Authors:** Fan Liu, Jing Yu, Yan-Xia Zhang, Fangzheng Li, Qi Liu, Yueyang Zhou, Shengshuo Huang, Houqin Fang, Zhuping Xiao, Lujian Liao, Jinyi Xu, Xin-Yan Wu, Fang Wu

**Affiliations:** Key Laboratory of Systems Biomedicine (Ministry of Education), Shanghai Center for Systems Biomedicine, Shanghai Jiao Tong University, Shanghai, 200240, China; State Key Laboratory of Microbial Metabolism, Sheng Yushou Center of Cell Biology and Immunology, School of Life Science and Biotechnology, Shanghai Jiao Tong University, Shanghai, 200240, China; School of Chemistry & Molecular Engineering, East China University of Science and Technology, Shanghai, 200237, China.; State Key Laboratory of Natural Medicines and Department of Medicinal Chemistry, China Pharmaceutical University, Nanjing, 210009, China; Hunan Engineering Laboratory for Analyse and Drugs Development of Ethnomedicine in Wuling Mountains, Jishou University, Hunan, 416000, China; Shanghai Key Laboratory of Regulatory Biology, School of Life Sciences, East China Normal University, Shanghai, 200241, China.

**Keywords:** Ammonia, High-throughput screening, Antibiotic resistance, Urease, Mechanism of action, *Helicobacter pylori*

## Abstract

To date, little attempt has been made to develop new treatments for *Helicobacter pylori* (*H. pylori*), although the community is aware of the shortage of treatments for *H. pylori*. In this study, we developed a 192-tandem-microwell-based high-throughput-assay for ammonia that is a known virulence factor of *H. pylori* and a product of urease. We could identify few drugs, i.e. panobinostat, dacinostat, ebselen, captan and disulfiram, to potently inhibit the activity of ureases from bacterial or plant species. These inhibitors suppress the activity of urease via substrate-competitive or covalent-allosteric mechanism, but all except captan prevent the antibiotic-resistant *H. pylori* strain from infecting human gastric cells, with a more pronounced effect than acetohydroxamic acid, a well-known urease inhibitor and clinically used drug for the treatment of bacterial infection. This study offers several bases for the development of new treatments for urease-containing pathogens and to study the mechanism responsible for the regulation of urease activity.

## 1. Introduction

Bacteria, fungi and plants, with the exception of animals, contain urease[1]. Urease (EC 3.5.1.5) is a class of nickel metalloenzyme that hydrolyzes amino acid metabolites to produce ammonia (NH_3_) and carbon dioxide[2, 3]. The active catalytic site of urease consists of two nickel ions, a carbamylated lysine residue, two histidines and an aspartic acid. In addition to the consistent catalytic mechanism, the amino acid sequence of urease has been reported to be highly conserved between different species[4].

Bacterial urease is known to be a key virulence factor of some pathogens for a number of diseases[5], e.g., *Helicobacter pylori* (*H. pylori*) for gastritis or gastric cancer, and *Proteus mirabilis* (*P. mirabilis*) for urinary tract infections and urinary stones[6]. The pathogens can hydrolyze urea substrates to produce NH_3_. The released NH_3_ not only helps *H. pylori* to survive in the low pH environment of the stomach but also causes damage to the gastric mucosa, triggering the infection[7]. Additionally, NH_3_ generated by *P. mirabilis* urease has been demonstrated to form urinary stones and destroy the urinary epithelium in the urinary system[8]. Because the human body does not contain urease, bacterial urease has been thought to be an important and specific drug target for combating these pathogens[9].

A number of studies have been performed to identify inhibitors of urease[10–13], but only one urease inhibitor, acetohydroxamic acid (AHA), was approved for the treatment of urinary infections and urinary stones in 1983 by the US Food and Drug Administration (FDA)[14, 15]. Severe side effects, low stability in gastric juice, and a lack of direct evidence for suppressing the growth of pathogens seem to be the limiting factors for the low success rate of these urease inhibitors. Adverse side effects of AHA, including teratogenic effects[16], a low efficiency indicated by the required high dose for the patient (∼ 1000 mg/day for adults), and the assumed drug resistance of bacteria, further imply that potent and bioactive inhibitors with new chemical moieties are urgently needed to combat these pathogens. Indeed, the current clinical first-line regimen for the treatment of *H. pylori* [proton-pump inhibitor, clarithromycin, amoxicillin or metronidazole (sometimes tinidazole)][17, 18], is unable to completely eradicate *H. pylori* due to the increased antibiotic resistance[17, 19].

To date, few validated high-throughput assay has been constructed to quantitatively analyze NH_3_ and the activity of NH_3_-generating enzyme urease, but no high-throughput screening approach has been employed to systematically extend the chemical moiety of urease inhibitors. The current assay to determine the activity of urease mainly relies on colorimetric reactions to determine the concentration of NH_3_ using indophenol or Nessler’s reaction[20]. Recently, a microfluidic chip-based fluorometric assay has been developed to monitor the activity of urease[21, 22]. In addition, a cell-based assay for *H. pylori* urease has been reported lately, and validated by known inhibitors of urease, but it has not been employed to screen new inhibitors for urease yet[23]. Overall, the current assay setting and procedures are relatively time-consuming and vulnerable to interference.

In this study, we established and validated a new tandem-well-based HTS assay for NH_3_ and NH_3_-generating urease and performed an HTS screening campaign to identify druggable chemical entities from 3,904 FDA or Foreign Approved Drugs (FAD)-approved drugs for jack bean and bacterial ureases. Five clinically used drugs, i.e., panobinostat, dacinostat, ebselen (EBS), captan and disulfiram, were found to be submicromolar inhibitors of *H. pylori* urease (HPU), jack bean urease (JBU), or urease from *Ochrobactrum anthropi* (*O. anthropi*), a newly identified pathogen with resistance to β-lactam antibiotics[24]. Moreover, panobinostat, dacinostat, EBS and disulfiram potently inhibited the infection of *H. pylori*, suggesting that these pharmacologically active moieties or drugs could serve as bases for the development of new treatments for urease-positive pathogens.

## 2. Material and methods

### 2.1 Materials

Jack bean urease (JBU), DMSO, and dithiothreitol (DTT) were purchased from Sigma (Steinheim, Germany). Hypochlorous acid, sodium nitroprusside, salicylate, potassium sodium tartrate, urea, sodium hydroxide, bovine serum albumin, Triton X-100, L-histidine and L-cysteine were purchased from Sangon (Shanghai, China). Nessler’s reagent was purchased from Jiumu company (Tianjin, China). Acetohydroxamic acid was purchased from Medchemexpress (Monmouth Junction, NJ). Columbia blood agar plate, liquid medium powder for *H. pylori*, bacteriostatic agent and polymyxin B were purchased from Hopebio company (Shandong, China). RMPI 1640 medium and fetal bovine serum (FBS) were purchased from Gibco (Invitrogen, Gaithersburg, MD). The other materials were purchased from the indicated commercial sources or were from Sigma.

### 2.2 Construction of the high-throughput screening assay for urease

The assay was constructed to measure the activity of urease based on a 192-tandem microwell plate, which we had previously developed to detect the H_2_S gas generated by H_2_S-generating enzymes[25, 26]. Phosphate or Tris buffer at various pH values were used to determine the optimal pH for JBU in the presence of 25 mM urea substrate (Figure S1E). The optimal conditions were found to be the 50 mM phosphate buffer and pH 7.4. Moreover, the suitable detection reagent and enzyme concentrations were resolved by testing three types of NH_3_ detection reagents with various concentrations of JBU or HPU, i.e., salicylic acid-hypochlorite, Nessler’s reagent and phenol red detection reagent (Figures S1A-C). The optimized conditions for the standard assay were found to be with salicylic acid-hypochlorite and commercial Nessler’s detection reagents (Jiumu, Tianjin, China) for JBU and HPU, respectively, in the presence of 50 nM JBU or 200-400 nM HPU, 25 mM urea, 100 μM NiCl_2_, and 50 mM phosphate buffer (final concentrations of pH 7.4). The salicylic acid-hypochlorite detection reagent contained 1.6 mM hypochlorite, 400 mM sodium hydroxide, 36 mM salicylic acid, 18 mM potassium sodium tartrate and 1.6 mM sodium nitroprusside. The assay was performed using multichannel pipettes to add 1 μl of each compound (solubilized in DMSO or H_2_O) and 24 μl of the enzyme mix (100 nM, 100 μM Tris, pH 7.9) into the reaction well (Figure 1A), followed by a 30-min incubation. After addition of 50 μl of salicylic acid-hypochlorite or Nessler’s detection reagent to the detection well, 25 μl substrate solution (50 mM urea, 200 μM NiCl_2_, 0.04% bovine serum albumin (w/v)) was mixed with the enzyme in the reaction well. The reaction was monitored at 37 °C, and the absorbance at 697 nm or 420 nm was accordingly measured at the appropriate time points in a microplate reader (Synergy2 from BioTek, Winooski, VT).

**Figure 1.**
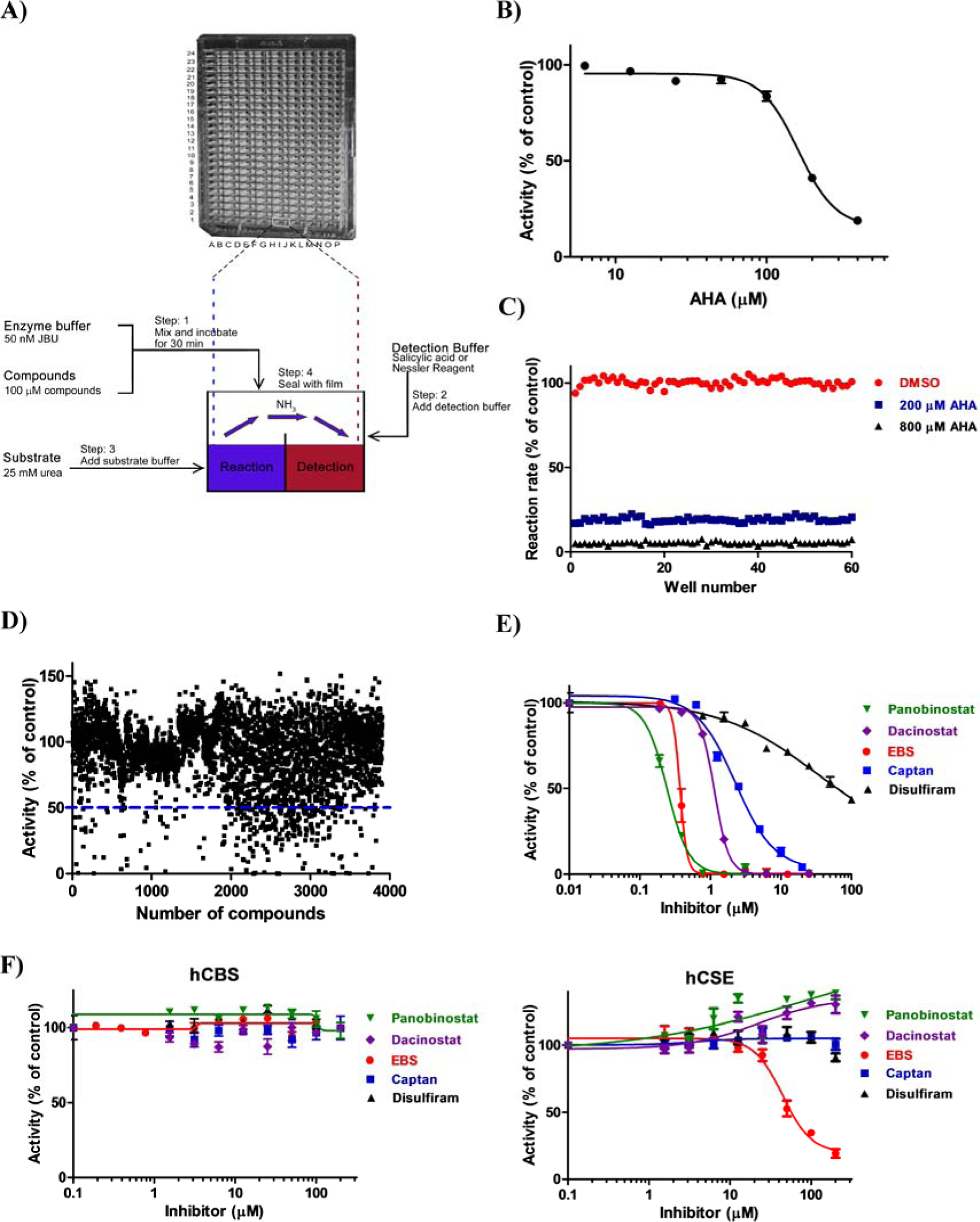
Development of a new high-throughput assay for urease and the discovery of new urease inhibitors. (**A**) Diagram of the tandem-well-based assay for the NH_3_-producing enzyme. The procedures for the assays and the cross-section of a tandem-well are shown. Blue, the reaction reagent; red, the detection reagent for NH_3_. (**B**) Validation of the urease assay with the known inhibitor AHA. (C) Well-to-well reproducibility of the 192-tandem-well-based assay for urease. ●, 2% DMSO (control, 100%); ▪, 200 μM AHA; ▲, 800 μM AHA (n = 60). **(D)** High-throughput inhibitor screening for JBU with 192-tandem-well plates. Compound concentration: 100 μM. **(E-F)** Dose-dependent effects of panobinostat, dacinostat, EBS, captan and disulfiram on the activity of JBU (**E**), human CBS (**F**) or human CSE (**F**). Means ± SDS (n = 3). All experiments except the primary screening (**D)** were independently repeated at least twice, and one representative result is presented.

### 2.3 Primary screening of urease inhibitors using a high-throughput assay

We screened 3,904 compounds of FDA or FAD-approved drugs from Johns Hopkins Clinical Compound Library (JHCCL, Baltimore, MD) or from TopScience Biotech Co. Ltd. (Shanghai, China) at 100 μM for the inhibition of JBU under standard assay conditions with salicylic acid-hypochlorite detection reagent as described above. The Z’ value of the screening assay was calculated from 60 negative samples (2% DMSO) and 60 positive samples (800 μM AHA) and found to be more than 0.9 [27], indicating the assay is an excellent assay. Routinely, 16 negative samples and 8 positive samples were used to determine the assay performance, and screening data with a minimum Z’ value of 0.5 were accepted.

Compounds that show more than 50% inhibition were selected for the further validation. Primary hits were defined as that compound is free of heavy metal atom and shows a more than 50% inhibition at 50 μM.

### 2.4 Compounds used for follow-up studies

All hits identified from the primary screening and their analogs were reordered in the highest pure powder from commercial sources or synthesized in-house for the following studies: dose-dependent, kinetic studies, biophysical assays, LC-MS/MS analysis, cell or bacteria-based studies. Panobinostat and dacinostat were brought from AdooQ (catalog number: A10518 for panobinostat, A10516 for dacinostat). EBS and captan were purchased from Sigma (catalog number: E3520 for EBS, 32054 for captan). Disulfiram (tetraethylthiuram disulfide) was purchased from TCI Chemicals (B0479). Captafol (1ST21228) was purchased from Alta Scientific Co.,Ltd (Tianjing, China), and dibenzyl diselenide (catalog number: B21278) was purchased from Alfa Aesar (Ward Hill, MA). Abexinostat (catalog number: HY-10990), belinostat (HY-10225), vorinostat (HY-10221), ricolinostat (HY-16026), ilomastat (HY-15768) and pracinostat (HY-13322) were brought from Medchemexpress. The purities of these commercially available primary leads or analogs of leads as well as in-house synthesized EBS derivatives were confirmed to be at least 95% by using HPLC (for details, see below), with an exception for EBS, the purity of which is determined with combustion analysis methods by the supplier. All the HPLC spectra as well as the combustion analysis data for these inhibitors, which were determined either from commercial supplier or by ourself, were included in the Supporting Information (see below).

### 2.5 Determination of IC_50_ values

The IC_50_ values of all the hits or their analogs, as well as AHA, on the activity of JBU, HPU or OAU were determined according to the above-described standard assay conditions. Compounds were incubated with the enzyme and assayed at a series of concentrations (at least 7 steps of doubling dilution). Similarly, the IC_50_ values of these inhibitors for hCBS or hCSE were determined accordingly[25]. Sigmoidal curves were fitted using the standard protocol provided in GraphPad Prism 5 (GraphPad Software, San Diego CA). IC_50_ was calculated by semilogarithmic graphing of the dose-response curves.

### 2.6 Aggregation-based assay

To exclude the mechanism by which inhibitors suppress the activity of urease via colloidal aggregation, we performed an aggregation-based assay in the presence of nonionic detergents[28]. Freshly prepared Triton X-100 (Sangon, Shanghai, China) at different concentrations of 0.1%, 0.05%, 0.01%, 0.005%, and 0.001% was first tested for its effects on the activity of JBU under standard assay conditions. Subsequently, the inhibitory effects of panobinostat, dacinostat, EBS, captan and disulfiram, as well as the analogs of EBS in the *in vitro* JBU activity assay, were determined in the presence of 0.01% Triton X-100, a concentration that alone has no inhibitory effect on the activity of JBU.

### 2.7 Reversibility assay

To illustrate the mode of action for the inhibitors of urease, we performed the rapid-dilution experiment. After incubation with panobinostat at a concentration of 4 μM, dacinostat at 10 μM, EBS or captan at 200, 100, 50 or 20 μM for 60 min, JBU (10 μM) was diluted 200-fold in the assay buffer. After a further incubation of 0, 1, 1.5, 2, 3, 4 or 5 h, the remaining activity of JBU was accordingly measured (METHODS). The inhibitor concentrations after dilution are indicated in the figure.

### 2.8 Determination of k_inact_ or K_I_ parameters for irreversible inhibitors

The IC_50_ values of EBS or captan for JBU were measured after different preincubation periods with the enzyme, i.e., 5, 10, 20, 30, 40, 45, 60, 70 or 90 min. The k_inact_ and K_I_ values for EBS or captan were obtained by nonlinear regression plotting of the time-dependent IC_50_ data as previously reported[29].

### 2.9 Enzyme kinetics

The reaction rate was determined with JBU at the indicated concentrations of panobinosta, dacinostat, EBS or captan against increasing concentrations of urea substrate (15.625, 31.25, 62.5, 125, 250, 500, 1000 mM for panobinosta and dacinostat; 12.5, 25, 50, 100, 200 mM for EBS and captan). The data were fitting to the Michaelis-Menten inhibition equation for determination of the competitive and noncompetitive inhibition parameter Ki and αKi using GraphPad Prism 5 (Table 1, Figures 2C and 3B)[25], respectively. To illustrate the inhibition type, Lineweaver-Burk plots of these inhibitors were drawn and analyzed.

**Figure 2.**
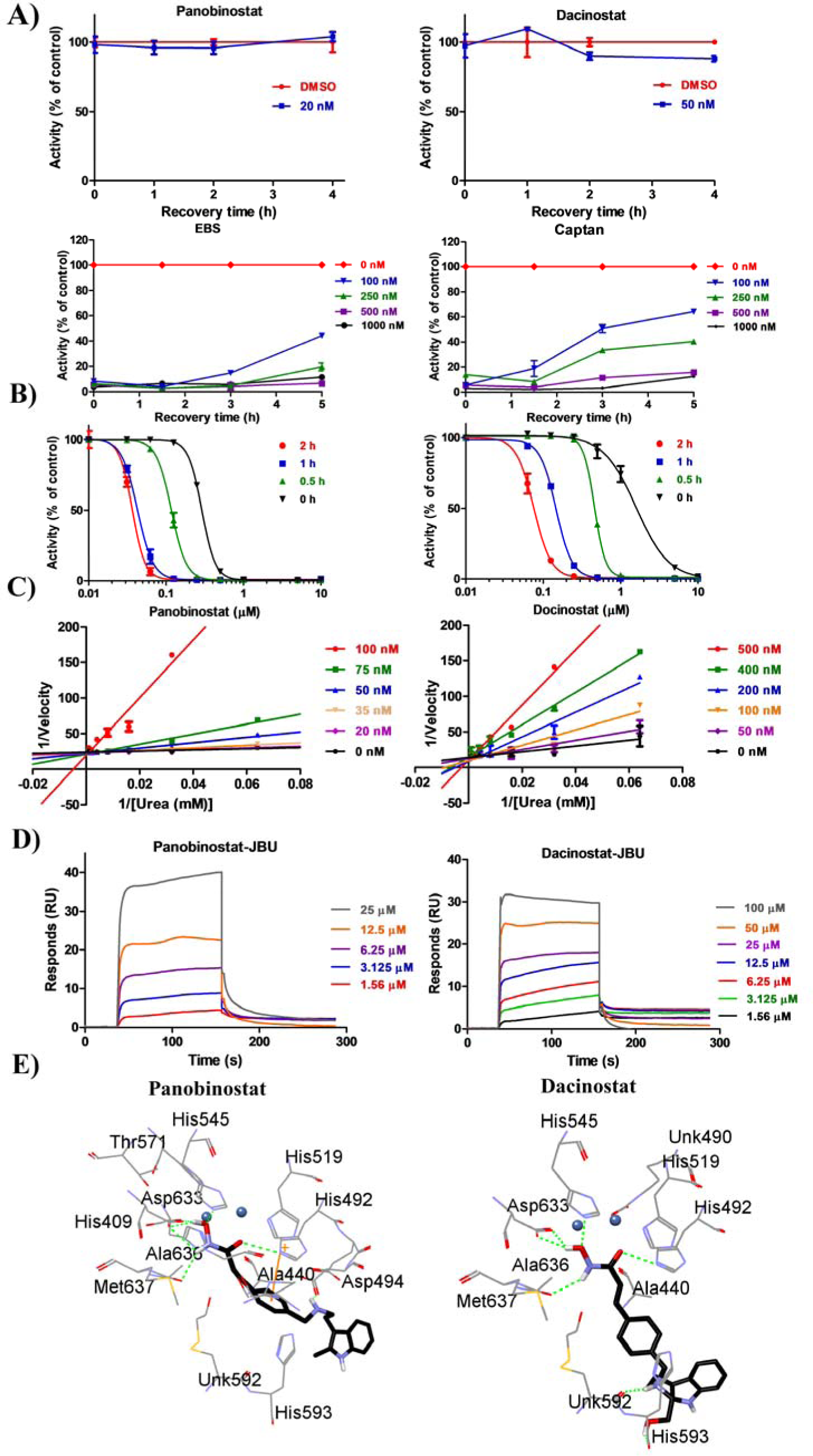
Panobinostat, dacinostat, EBS and captan inhibit the activity of JBU. (**A**) Panobinostat and dacinostat are reversible inhibitors, whereas EBS and captan are covalent inhibitors or slow-binding inhibitors toward JBU. Means ± SDs (n = 3). (**B**) Effects of the incubation period on the IC_50_ values of panobinostat and dacinostat toward JBU. Panobinostat and dacinostat were preincubated with JBU for the indicated times before performing the standard assay to analyze their inhibitory effects. Means ± SDs (n = 3). (**C**) Inhibition of JBU by panobinostat or dacinostat as a function of urea concentration. K_i_ values for panobinostat and dacinostat, 0.02 μM and 0.07 μM, respectively. Means ± SDs (n=3). (**D**) Surface plasmon resonance assay analysis of the binding of panobinostat or dacinostat to JBU. K_D_ were calculated using Biacore evaluation software and listed in Table 1. (**E**) The putative binding mode of panobinostat or dacinostat in the JBU active site. Panobinostat and dacinostat were docked into the JBU crystal structure (PDB code: 4GOA) using the Discovery Studio software. Residues surrounding the inhibitor within a distance of 3.5 Å are shown in gray; and hydrogen bonds are represented as green dotted lines. The experiments were independently repeated at least twice, and one representative result is presented.

**Figure 3.**
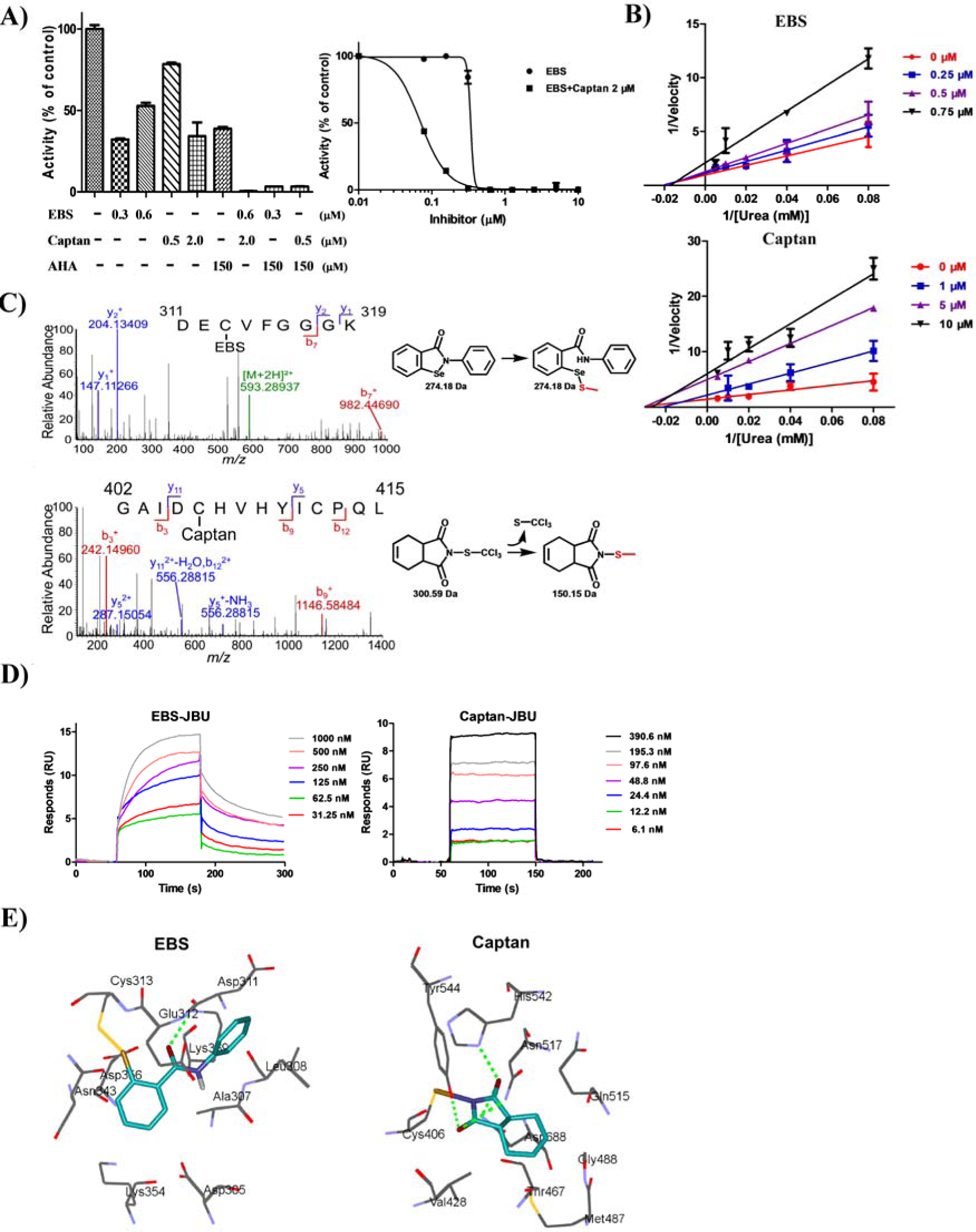
EBS or captan allosterically inhibits the activity of urease by covalently modifying a non-active-site Cys residue. (**A**) The synergistic inhibitory effects of the combinations of EBS, captan or AHA. A dose-dependent synergistic effect of the combination of EBS at the indicated concentrations with 2 μM captan was observed (right panel). Data are presented as percentages of the controls (DMSO and 2 μM captan alone in the left panel and right panel, respectively, 100%). Means ± SDs (n=3). (**B**) Inhibition of JBU by EBS or captan as a function of the urea concentration. αKi for EBS and captan, 0.8 μM and 1.1 μM, respectively. Means ± SDs (n=3). (**C**) Tandem mass spectrometry analysis of the modification site of EBS and captan on JBU. The Cys modification of EBS and captan on JBU were illustrated in the right panels. (**D**) Surface plasmon resonance assay analysis of the binding of EBS or captan to JBU. (**E**) The potential binding modes of EBS and captan in JBU. EBS and captan were modeled into the respective allosteric sites presented in the crystal structure of JBU (PDB code: 4GOA; METHODS). The residues within 3.5 Å surrounding the EBS and captan are shown. Hydrogen bonds are indicated as dashed green lines. The experiments were independently repeated at least twice, and one representative result is presented.

**Table 1.**
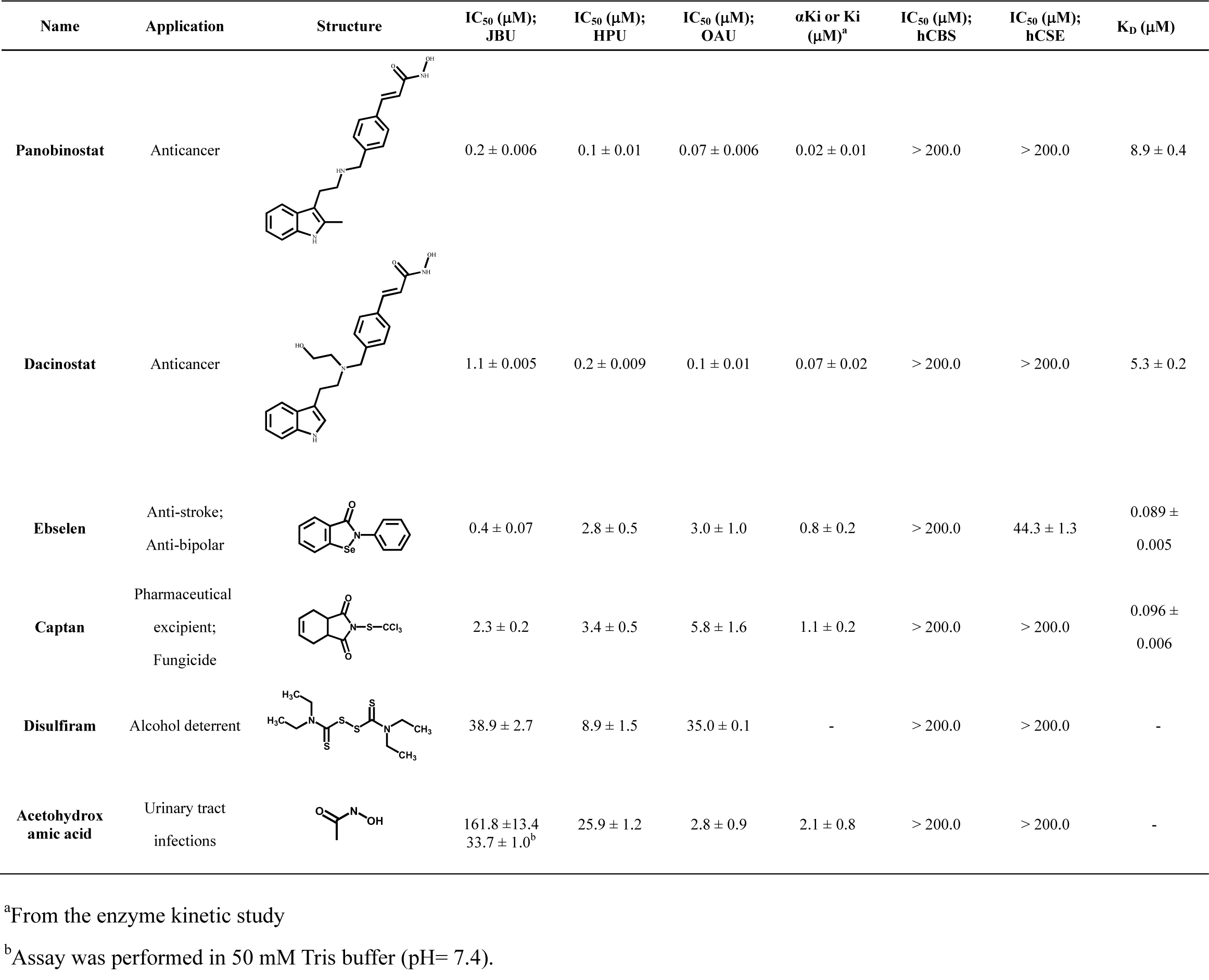
Indication, chemical structure, IC_50_, αKi, or K_D_ values of urease inhibitors.

| Name | Application | Structure | IC<sub>50</sub> (μM);<br>JBU | IC<sub>50</sub> (μM);<br>HPU | IC<sub>50</sub> (μM);<br>OAU | αKi or Ki<br>(μM)<sup>a</sup> | IC<sub>50</sub> (μM);<br>hCBS | IC<sub>50</sub> (μM);<br>hCSE | K<sub>D</sub> (μM) |
| --- | --- | --- | --- | --- | --- | --- | --- | --- | --- |
| Panobinostat | Anticancer |  | 0.2 ± 0.006 | 0.1 ± 0.01 | 0.07 ± 0.006 | 0.02 ± 0.01 | > 200.0 | > 200.0 | 8.9 ± 0.4 |
| Dacinostat | Anticancer |  | 1.1 ± 0.005 | 0.2 ± 0.009 | 0.1 ± 0.01 | 0.07 ± 0.02 | > 200.0 | > 200.0 | 5.3 ± 0.2 |
| Ebselen | Anti-stroke;<br>Anti-bipolar |  | 0.4 ± 0.07 | 2.8 ± 0.5 | 3.0 ± 1.0 | 0.8 ± 0.2 | > 200.0 | 44.3 ± 1.3 | 0.089 ±<br>0.005 |
| Captan | Pharmaceutical<br>excipient;<br>Fungicide |  | 2.3 ± 0.2 | 3.4 ± 0.5 | 5.8 ± 1.6 | 1.1 ± 0.2 | > 200.0 | > 200.0 | 0.096 ±<br>0.006 |
| Disulfiram | Alcohol deterrent |  | 38.9 ± 2.7 | 8.9 ± 1.5 | 35.0 ± 0.1 | - | > 200.0 | > 200.0 | - |
| Acetohydroxamic acid | Urinary tract<br>infections |  | 161.8 ± 13.4<br>33.7 ± 1.0<sup>b</sup> | 25.9 ± 1.2 | 2.8 ± 0.9 | 2.1 ± 0.8 | > 200.0 | > 200.0 | - |

<sup>a</sup>From the enzyme kinetic study
<sup>b</sup>Assay was performed in 50 mM Tris buffer (pH= 7.4).

### 2.10 LC-MS/MS analysis

JBU at a concentration of 12.5 μM was incubated with DMSO, 200 μM EBS or 200 μM captan for 120 min at room temperature. Then, three aliquots of 25 μg samples from the inhibitor-treated JBU or purified HPU (fraction 3 in Figure 4B) were digested separately with three proteases, including 0.5 μl trypsin (1 μg/μl), 0.5 μl GluC (1 μg/μl) or 0.5 μl subtilisin (1 μg/μl) overnight. The proteolytic peptides were combined and desalted on C18 spin columns and dissolved in buffer A (0.1% formic acid in water) for LC-MS/MS analysis. The peptides were separated on a 15-cm C18 reverse-phase column (75 μm × 360 μm) at a flow rate of 300 nl/min, with a 75-min linear gradient of buffer B (0.1% formic acid in acetonitrile) from 2% to 60%. The MS/MS analysis was performed on the Q-Exactive Orbitrap mass spectrometer (Thermo Fisher Scientific, San Jose, CA) using standard data acquisition parameters as described previously[30]. The mass spectral raw files were searched against the protein database derived from the standard sequence of JBU, HPU or the proteome of *H. pylori* using Proteome Discovery 1.4 software (Thermo Fisher Scientific, San Jose, CA), with a differential modification of 274.18 m/z in the case of EBS and 150.15 m/z in the case of captan.

**Figure 4.**
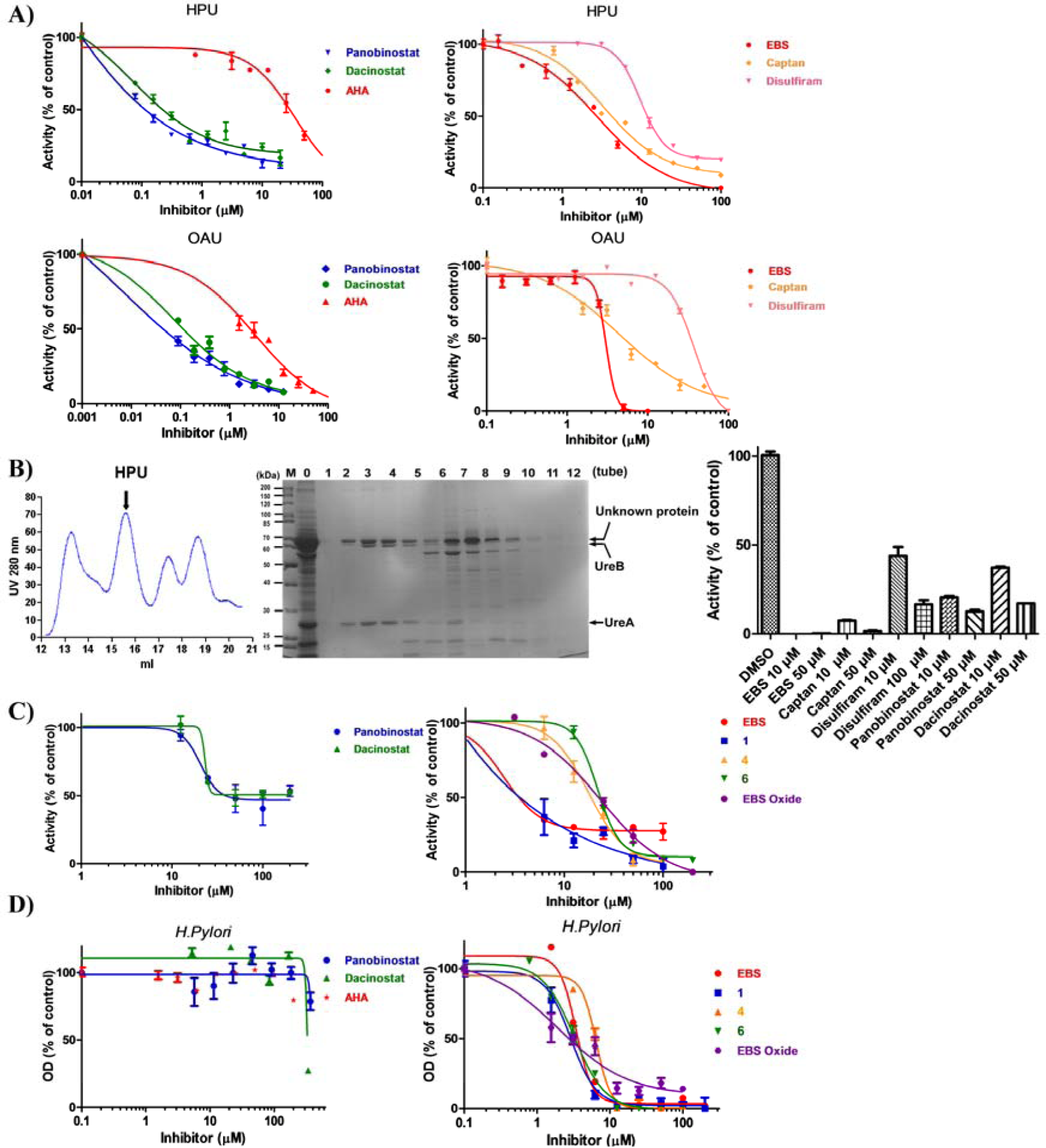
Urease inhibitors suppress bacterial ureases or the growth of urease-containing bacteria. (**A**) Dose-dependent effects of panobinostat, dacinostat, EBS, captan, disulfiram and AHA on the activity of *H. pylori* urease (HPU, upper panel) or *O. anthropic* urease (OAU, lower panel) *in vitro*. (**B**) Panobinostat, dacinostat, EBS, captan and disulfiram inhibit the activity of purified HPU from size-exclusion chromatography. Chromatography of the purification is shown in the left panel. The collected fractions (numbers 1-12) of the peaks (left panel), as well as the crude extract (number 0), were separated by 10% SDS-PAGE and stained with Coomassie Brilliant Blue R-250 (middle panel). The arrows indicate the peak of *H. pylori* urease (left panel) or subunit A or B of *H. pylori* urease (middle panel). The collected sample containing the urease (number 3) was tested to evaluate the inhibitory effects of indicated compounds (right panel). The protein identity of fraction 3 was analyzed by LC-MS/MS (METHODS and Figure S5). (**C**) The inhibitory effects of panobinostat, dacinostat and newly synthesized EBS analogs (**1**, **4** and **6**) on the activity of HPU in culture. Inhibitors were incubated with the *H. pylori* bacteria for 6 h. (**D**) The effects of panobinostat, dacinostat, EBS and its derivatives on the growth of *H. pylori.* Mean ± SD (n=3). All experiments were independently repeated at least twice, and one representative result is presented.

### 2.11 Surface plasmon resonance assays

The direct interactions between panobinostat, dacinostat, ebselen or captan and JBU were observed by the surface plasmon resonance (SPR) experiment with a BIAcore T200 (GE Healthcare, Uppsala, Sweden). JBU was immobilized on the surface of the CM5 sensor chip via the amino-coupling kit. The working solution used for the SPR assay was PBS-P (10 mM Na_2_HPO_4_, 1.8 mM KH_2_PO_4_, 2.7 mM KCl, and 140 mM NaCl in presence of 5% DMSO, pH 7.4). To determine the affinity of the inhibitors toward JBU, panobinostat, dacinostat, EBS or captan were diluted to specific concentrations with PBS-P buffer (for panobinostat: 25, 12.5 6.25, 3.125, 1.56 μM; dacinostat: 100, 50, 25, 12.5 6.25, 3.125, 1.56 μM; EBS: 1000, 500, 250, 125, 62.5, or 31.25 nM; for captan: 390.6, 195.3, 97.6, 48.8, 24.4, 12.2 or 6.1 nM) and subjected to the JBU-coated chips. The K_D_ values were calculated with BIAcore evaluation software (version 3.1).

### 2.12 Molecular modeling

The crystal structures of ureases were obtained from the Protein Data Bank (PDB code: 4GOA for JBU; PDB code: 1E9Y, HPU). The binding modes of panobinostat or dacinostat were gathered by using the CDOCKER module of the Discovery Studio software (version 3.5; Accelrys, San Diego, CA). Alternatively, AutoDock Vina was initially used to dock the EBS or captan to the respective Cys-containing allosteric site of JBU to obtain the appropriate configurations, enabling the reactive motifs of the compounds (the Se-containing benzisoxazole of EBS and the isoindole dione moiety of captan) to fall into the distance restraint of one covalent bond to the sulfur atom of the reactive Cys residue. The Se-S bond or the N-S bond for isoindole dione was then manually incorporated using the Discovery Studio 3.5 software (Accelrys, San Diego, CA). Subsequently, molecular dynamics simulation was performed with AMBER14 software and the ff03.r1 force field[31]. To relieve any steric clash in the solvated system, initial minimization with the frozen macromolecule was performed using 500-step steepest descent minimization and 2,000-step conjugate gradient minimization. Next, the whole system was followed by 1,000-step steepest descent minimization and 19,000-step conjugate gradient minimization. After these minimizations, 400-ps heating and 200-ps equilibration periods were performed in the NVT ensemble at 310 K. Finally, the 100-ns production runs were simulated in the NPT ensemble at 310 K. The binding modes for these inhibitors were visually inspected and the best docking mode was selected.

### 2.13 Bacterial strains and culture conditions

Bacterial strains of *H. pylori* or *O. anthropic* were obtained from BeiNuo Life Science (Shanghai, China). The strains were maintained on Columbia blood agar plates (Hopebio, Shandong, China) containing 5% defibrinated sheep blood at 37 °C under microaerobic conditions (5% O_2_, 10% CO_2_ and 85% N_2_), which was supplied by an AnaeroPack-MicroAero gas generator (Mitsubishi Gas Chemical Company, Japan). After a culture of 3-5 days in the plate, the bacterial colonies were scratched into the liquid medium for *H. pylori*, containing 10% or 7% fetal bovine serum and an antibacterial cocktail (composed of 10 mg/l nalidixic acid, 3 mg/l vancomycin, 2 mg/l amphotericin B, 5 mg/l trimethoprim and 2.5 mg/l polymyxin B sulfate; BeiNuo, Shanghai, China), and microaerobically incubated for another 3 or 5 days. Then, the medium or bacterial cells were collected for subsequent experiments.

A single colony of *O. anthropic* was inoculated into Luria-Bertani liquid medium (LB), which was supplemented with 50 mg/l ampicillin, 30 mg/l kanamycin and 10% FBS (Invitrogen) and cultured at 37 °C. After the bacterial culture reached an O.D. of 0.8 at 600 nm, the bacterial cells were collected by centrifugation for future experiments.

The identification of *H. pylori* and *O. anthropic* strain was carried out by PCR amplification of the urease gene or 16S rRNA with known primers (Table S4), LC-MS/MS analysis of proteins in the extracts, the bacterial urease activity assay or Gram staining.

### 2.14 16S rRNA sequencing

One colony from the *H. pylori* or *O. anthropic* culture plate was suspended in 50 μl of sterile water, and the DNA was liberated by a boiling-freezing method. The 16S rRNA gene was selectively amplified from this crude lysate by PCR using the universal primers 27f and 1492r, which have been previously described (Table S4). The PCR products at ∼1400 bp were sequenced. The resultant 16S rRNA sequences were compared with the standard nucleotide sequences deposited in GenBank with the BLAST program (http://www.ncbi.nlm.nih.gov/blast/). The DNA sequences of 16S rRNA extracted from these strains were confirmed to be from *H. pylori* or *O. anthropic*.

### 2.15 Preparation of crude extracts from the *H. pylori* and *O. anthropic* strains for the urease activity assay

For the urease activity assay, *H. pylori* or *O. anthropic* was cultured accordingly in 100 ml of broth medium as described above. Bacteria were centrifuged at 5,000 rpm for 30 min, and the pellet was washed with phosphate-buffered saline (PBS, pH = 7.4). The pellet was resuspended in 7 ml of PBS in the presence of protease inhibitors (Sigma-Aldrich, Steinheim, Germany) and then sonicated for 30 min of 30 cycles (30 s run and 30 s rest) using the noncontact ultrasonic rupture device (Diagenode, Liege, Belgium). The resultant bacterial lysate was centrifuged twice at 12,000 rpm for 30 min; the supernatant was collected and desalted using a Sephadex G-25 desalting column (Yeli, Shanghai, China). The protein in the fractions was separated by 10% SDS-PAGE, and the corresponding protein band for urease was quantified to determine the concentration of ureases by Coomassie blue R-250 (Sinopharm, Shanghai, China) staining using bovine serum albumin as a standard. The desalted fractions were stored at -80 °C in the presence of 15% glycerol until usage in the activity assay.

### 2.16 Size-exclusion chromatography for the purification of urease from *H. pylori*

The crude extract from *H. pylori* was first centrifuged at 12,000 rpm for 30 min. One milliliter of supernatant was loaded onto a gel filtration column (10 mm × 30 cm; GE Healthcare) and eluted with PBS at a rate of 0.5 ml/min on an AKTA Explorer 100 FPLC Workstation (GE Healthcare). The protein peaks observed were collected in Eppendorf tubes in a volume between 0.5 and 1 ml. The collected fractions were separated by PAGE on a 10% Tris-glycine SDS-gel and stained with Coomassie Brilliant Blue R-250 to identify *H. pylori* urease.

### 2.17 Determination of the minimal inhibition concentration and dose-dependent growth-inhibition curve for urease inhibitors

The minimal inhibition concentration (MIC) and dose-dependent growth-inhibition curve for the inhibitors on *H. pylori* were determined using the broth dilution method[32]. Briefly, *H. pylori* was grown to an OD_600_ nm of 1.0 in liquid medium supplemented with 7% FBS under standard culture conditions. Then, 150 μl *H. pylori* in the diluted culture (OD of 0.1) was incubated with the inhibitors at final concentrations of 1, 2, 4, 16, 32, 64, 128, 256, 512 μg/ml or at indicated concentrations for 72 h. The OD_600_ nm was measured to calculate the percentage of growth inhibition. The DMSO (1% final concentration)-treated *H. pylori* cultures and culture medium in the absence of bacteria were referred as the negative control (0%) and positive control (100%), respectively. The MIC was defined as the lowest concentration of inhibitor that inhibited 100% of bacterial growth. The *H. pylori* strain was found to be resistant to tinidazole or metronidazole and have an MIC of greater than 512 μg/ml.

### 2.18 Bacterial-cell-based assay for measuring the activity of urease in culture

The endogenous activity of HPU in bacterial cultures was determined using the tandem-well-based plate. Briefly, 300 μl of *H. pylori* culture (OD_600 nm_ ∼1.0) was treated with panobinostat, dacinostat or EBS as well as EBS analogs for 6 or 24 h at different concentrations (0, 3.125, 6.25, 12.5, 25, 50, 100 or 200 μM). Then, the bacterial cells were centrifuged, washed and resuspended in assay buffer containing 25 mM urea. Finally, the ∼100 μl suspension was added to the reaction well of the tandem-well plate and assessed for the activity of urease with Nessler’s reagent under standard assay conditions.

### 2.19 Gastric cell infection model of *H. pylori*

The cell infection model of *H. pylori* was constructed using the SGC-7901 adenocarcinoma gastric cell line and following an established protocol[16]. Briefly, *H. pylori* was cultured in liquid medium for *H. pylori* at 37 °C for 3-5 days under standard culture conditions (see above). Then, *H. pylori* at a concentration of 1.5 × 10^6^ CFU/ml was treated with the indicated inhibitors for 24 h in culture. The bacterial suspension together with 10 mM urea were subsequently added to the culture medium of SGC-7901 cells (MOI = 30), which had been cultured with RPMI 1640 medium plus 10% FBS in a 96-well plate for one day, and coincubated with the cells for an additional 24 h. Cell images were obtained at specific time points prior to and one day after addition of the bacterial culture using Image Xpress Micro^®^ XLS (Molecular Devices, Sunnyvale, CA) under a 20 × objective lens. The cell numbers in the images were quantified using Image Xpress Software. The protective effects of the inhibitors were calculated by dividing the number of SGC-7901 cells after the 24-h treatment by that prior to the treatment (100%) in the same well.

## 3. Results

### 3.1 Development of a high-throughput assay and identification of potent inhibitors for urease

To construct a high-throughput assay for NH_3_-generating urease and prevent the detection interference from substances in the enzyme extraction, we utilized a 192-tandem-well-based gas-detection method, which we previously developed to monitor the activity of H_2_S-generating enzymes[25, 26]. The tandem-well design could physically separate the gas product from the enzymatic reaction and enable the specific and real-time detection of the gas-producing enzyme activity (Figure 1A).

To construct the HTS assay, we compared three reported protocols for determination of the activity of JBU by using salicylic acid-hypochlorite and Nessler detection reagent, as well as phenol red[21, 33, 34], which undergo the indophenol and Nessler’s reaction with NH_3_, respectively. Salicylic acid-hypochlorite and Nessler’s reagents could dose-dependently and time-dependently monitor the activity of JBU at various concentrations (Figures S1A and B); however, the phenol red failed to detect it (Figure S1C). We decided to choose salicylic acid-hypochlorite as the detection reagent for the HTS screening assay of JBU (Figure S1A) due to its lower toxicity than Nessler reagent, which contains mercury[33]. The absorbance (OD) at 697 nm of the blue complex indophenol generated from salicylic acid was correlated linearly with the concentration of NH_4_Cl (19.5 - 625 μM), thus validating the analytic setup for NH_3_ quantification (Figure S1D). Moreover, the optimal assay buffer for JBU was found to be phosphate buffer at pH 7.4 (Figure S1E). In contrast, we employed Nessler’s reagent to detect the activity of HPU and *Ochrobactrum anthropic* urease (OAU) in subsequent studies since it showed a better sensitivity for the limitation of detection of the activity of HPU than salicylic acid-hypochlorite (Figures S1F and 1G). Collectively, we chose 50 nM of JBU and 25 mM urea substrate in the phosphate buffer to perform the assay.

Under the assay conditions, AHA showed an IC_50_ of ∼ 160 μM (Figure 1B), which was very similar to the previously reported value (IC_50_ of ∼ 140 μM; ref. [14]), indicating that the newly developed assay for urease was accurate and reliable. However, the IC_50_ of AHA was found to decrease to 33.7 μM when using the 50 mM Tris buffer instead of the phosphate buffer in our assay (Table 1). To determine the well-to-well reproducibility, the assay was validated with 200 μM AHA (∼ IC_50_) or 800 μM (∼ 5-fold IC_50_) AHA. The tandem-well plate consistently showed distinct differences among the control, the 200 μM-AHA-treated and the 800 μM-AHA-treated groups (Figure 1C). The average Z’ values of the assay were found to be ∼ 0.9 when they were calculated with the 800 μM AHA positive control.

To identify novel and potent inhibitors for urease, we screened 3,904 FDA or FAD-approved drugs at 100 μM. Five potent hits, i.e., panobinostat, dacinostat, EBS, captan and disulfiram, were found to dose-dependently inhibit the activity of JBU with IC_50_ values of 0.2, 1.1, 0.4, 2.3 and 38.9 μM, which are ∼ 800, 146, 400, 70, 4, -fold more potent than AHA, respectively (Figure 1E and Table 1). Intriguingly, the former two drugs are analogs of AHA. Importantly, all of them seemed to bear significant selectivities for urease since they did not substantially inhibit other gas-producing enzymes, i.e., cystathionine beta-synthase (CBS) and cystathionine γ-lyase (CSE), two H_2_S-generating enzymes (Figure 1F). Moreover, the potent inhibitory effects of these inhibitors were likely due to on-target inhibition of JBU rather than the nonspecific reaction with NH_3_ or forming an aggregation since they did not react with NH_3_ and their inhibition was not attenuated by the detergent (Figures S2A and S2B). In corroborating these findings, EBS and disulfiram have recently been reported to be specific inhibitors of bacterial and plant urease[11, 12], respectively, although their mode of actions for inhibiting urease, and their effects on the proliferation or infection of urease-containing pathogens remain little explored.

### 3.2 The mode of action study for urease inhibitors

To determine the reversibility of the inhibition by panobinostat, dacinostat, EBS, captan and disulfiram to JBU, various concentrations of the inhibitors and JBU were incubated together for 60 min (Figure 2A). After a 200-fold dilution, the inhibitory effects of panobinostat and dacinostat as well as disulfiram were found to be reversible (Figures 2A and S3C). In contrast, EBS or captan at 100 nM was found to completely block the activity of JBU; this concentration did not affect the activity without the pre-incubation with enzyme (Figure 1E). Additionally, the inhibitions exerted by EBS or captan were not fully recovered (Figure 2A), indicating that both of them were likely to be covalent or slow-dissociation inhibitors for JBU.

Surprisingly, the inhibitory effect of disulfiram was found to be dependent on the concentrations of Ni^2+^ ion, the catalytic cofactor for urease (Figure S3C), indicating that it inhibits JBU likely via formation of a complex with the catalytic Ni^2+^ ion and subsequently occupying the active site of JBU. This explanation seems to be plausible since recent findings have revealed that disulfiram inhibits the proliferation of tumor cells by forming a complex with Cu^2+^[35].

Moreover, the inhibitory potencies of panobinostat and dacinostat were found to increase with the pre-incubation time of the compound with urease (Figure 2B). After 2 h pre-incubation, the IC_50_ value of panobinostat and dacinostat were decreased ∼ 7.5 folds and ∼ 18.8 folds, respectively (Figure 2B). In enzyme kinetics studies for JBU, panobinostat and dacinostat were found to be competitive inhibitors towards urea substrate, with a K_i_ value of 0.02 and 0.07 μM (Figure 2C and Table 1), which are ∼ 105 folds and 30 folds more potent than AHA (K_i_ ∼ 2.1 μM; Table 1). However, the inhibition of these two inhibitors was not dramatically interfered with the supplemented Ni^2+^ (Figure S3A). Also, the addition of histidine or cysteine has no effects on the inhibition of panobinostat or dacinostat (Figure S3B). Importantly, the surface plasmon resonance assay demonstrate that these two compounds could physically bind to JBU (Figure 2D; Table 1). The drastic effect seems not only relying on the hydroxamic acid motif that is the known pharmacophore of AHA-derivative inhibitors, but also the hydrophobic ring and secondary amine group, as indicated by that the benzene ring favorably interacts with the His492 residue and/or the nitrogen atom forms an additional hydrogen bond with Asp494 in the modeled inhibitor-JBU complex structure (Figure 2E).

In contrast, the inhibition caused by both EBS and captan was found to be prevented by the addition of dithiothreitol (DTT) or free cysteine into the enzymatic reaction, but not that of histidine or Ni^2+^ (Figures S4A-C). Furthermore, the IC_50_ values of the two inhibitors were linear with the concentrations of the enzyme (Figure S4D), an inhibitory feature of the covalent inhibitor[29], confirming that they targeted the enzyme covalently. The inhibition constants for these irreversible inhibitors, i.e., the rate of enzyme inactivation (k_inact_) and inactivation rate constants (K_I_), were also determined by nonlinear regression of the time-dependent IC_50_ values (Figure S4E) [29]. The k_inact_ and K_I_ for EBS were found to be 2.79 × 10^-3^ s^-1^ and 0.73 μM, which were 4.4 and 2.4-fold better than captan (k_inact_, 0.63 × 10^-3^ s^-1^; K_I_, 1.76 μM), respectively. Taken together, the results demonstrated that EBS and captan inhibited JBU by covalently modifying the Cys rather than His residue, the latter of which is known to be the active site of urease [2, 3]. Interestingly, we observed a synergistic inhibitory effect from the combination of EBS and AHA (Figure 3A), a substrate-competitive inhibitor for urease, implying that EBS targeted Cys residue(s) of another site rather than the active site. Similar experimental results were also obtained for captan. Moreover, the combination of EBS with 2 μM captan also significantly increased the potency of EBS by 6-fold (right panel, Figure 3A), implying distinct binding sites of the two covalent inhibitors.

To corroborate this finding, we performed enzyme kinetics, mass spectrometry and surface plasmon resonance studies (Figures 3B-D). Consistently, EBS or captan displayed a noncompetitive mode for the urea substrate (Figure 3B). Furthermore, tandem-mass spectrometry analysis revealed that Cys313 and Cys406, which were not adjacent to the active site, appeared to be modified by EBS and captan, respectively (Figure 3C). The addition of 274.18 daltons in molecular weight was observed for EBS, demonstrating the breakage of the Se-N bond and formation of the Se-S bond with the Cys residue, a phenomenon that has been reported previously for EBS[36]. However, the increase of 150.15 daltons suggested that only the isoindole dione moiety of captan modified the Cys residue, accompanied by the release of the trichloromethyl thio moiety [-SC(C1)_3_]. This new observation provides a new perspective for the unexplored covalent molecular mechanism of captan.

Additionally, a potent and physical interaction between EBS or captan and JBU was observed in the surface plasmon resonance study (Figure 3D). The equilibrium dissociation constant (K_D_) for EBS and captan was found to be 89 and 96 nM, respectively.

To illustrate the binding mode of EBS or captan, we modeled them into the respective allosteric Cys-containing pocket (Cys313 for EBS, Cys406 for Captan) in JBU by using molecular dynamics simulations (Figure 3E). The carbonyl group of EBS was found to form a hydrogen bond with Lys369, and the phenyl ring interacts with the hydrophobic side chain of Leu308. Additionally, the two carbonyl groups of captan formed four hydrogen bonds with the side chains of Asn517, His542, Tyr544 and Asn688. Taken together, these results implied that these intermolecular weak interactions also substantially contributed to the binding of the covalent inhibitors to the protein, in addition to the covalent interaction.

### 3.3 The inhibitory effect of inhibitors on bacterial ureases

Next, we investigated the effects of panobinostat, dacinostat, EBS and captan as well as disulfiram on the activity of HPU and OAU, two bacterial ureases from *H. pylori* and *O. anthropic*, respectively. As expected, these drugs could inhibit the activity of HPU in the crude extracts and showed IC_50_ values of 0.1 μM, 0.2 μM, 2.8 μM, 3.4 and 8.9 μM, which indicated that they were ∼ 259, 130, 10, 8 and 3 -fold more potent than AHA (IC_50_ ∼ 25.9 μM; Figure 4A and Table 1), respectively. Moreover, panobinostat, dacinostat, EBS, captan and disulfiram were also found to inhibit the partially purified HPU, which was isolated by size-exclusion chromatography (Figures 4B and S5). Consistently, they also suppressed the activity of OAU at a similar potency to HPU (Figure 4A and Table 1). Compounds **1, 4** and **6**, which were synthesized in house (Scheme S1), as well as commercially available EBS oxide, also showed a better efficiency than EBS (IC_50_ ∼ 2.8 μM) in the *in vitro* HPU-based enzyme assay (Table S1), and **4** displayed a maximum three-fold increase in potency (IC_50_ ∼ 1.1 μM; Table S1). Moreover, we could confirm that panobinostat, dacinostat and EBS as well as EBS oxide, **1**, **4** or **6,** could largely suppress the activity of HPU in culture (Figure 4C). The IC_50_ values of these inhibitors for inhibiting the urease of the cultured *H. pylori* strain ranged from 5.7 to 23.2 μM (Figure 4C and Table S2).

**Figure 5.**
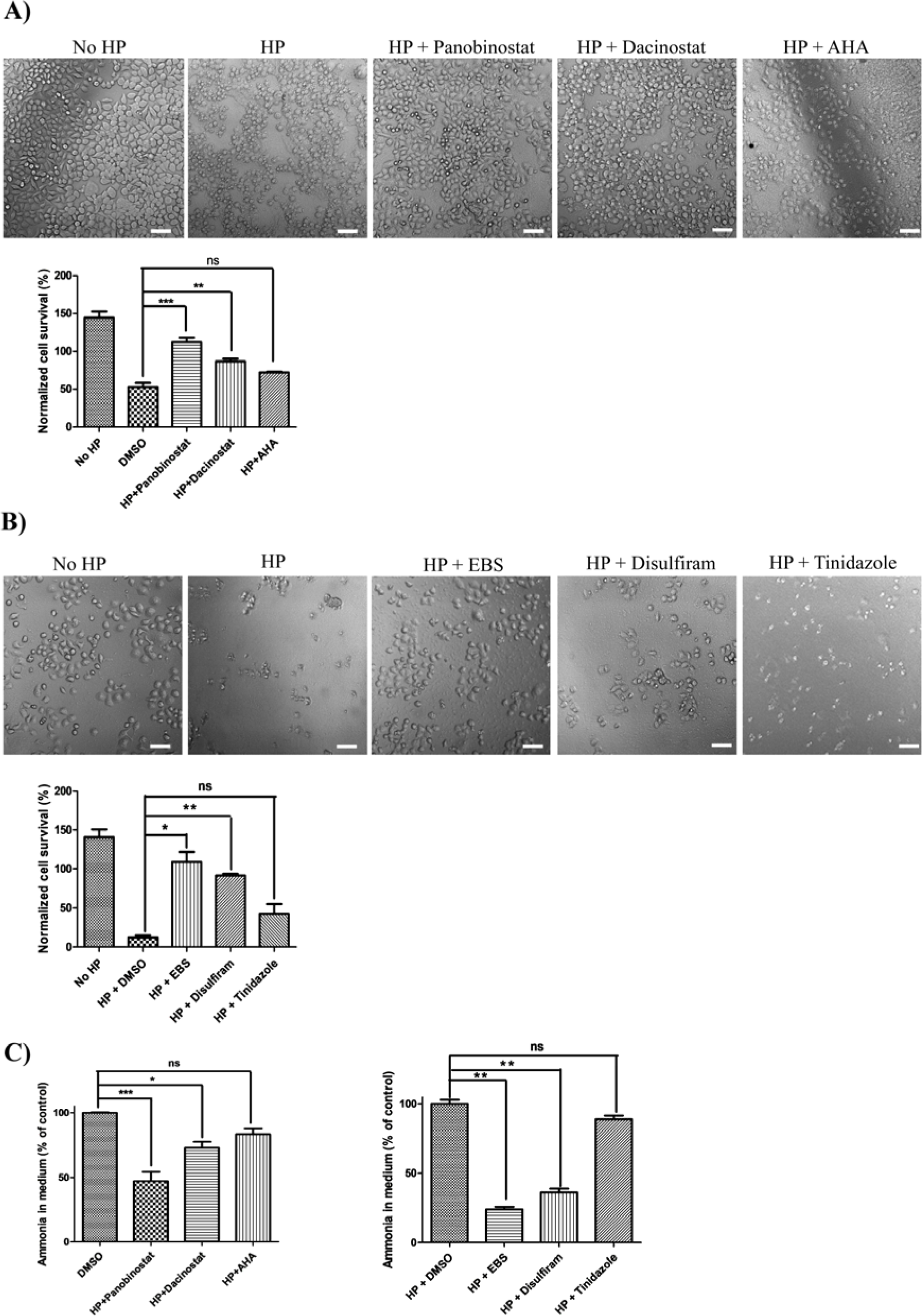
Panobinostat, dacinostat and EBS inhibits the virulence of *H. pylori* in cultured gastric cells. SGC-7901 cells were infected with HP in the presence of 30 μM panobinostat (**A**), 30 μM dacinostat (**A**), 30 μM AHA (**A**), 20 μM EBS (**B**), 20 μM disulfiram or 50 μM tinidazole (**B**) for 24 h before capturing the images in bright field by Image Xpress Micro® XLS (Molecular Devices, Sunnyvale, CA) under a 20 × objective lens. A representative image for each treatment condition is shown (n = 3). Scale bars, 100 μm. The cell numbers before treatment (100%) or after 24 h of treatment were quantified. (**C**) The effects of urease inhibitors on the NH_3_ amount of the cell culture medium. After the treatment, the amount of NH_3_ in the cell medium of the corresponding samples was quantified with Nessler’s reagent, and the data are shown as percentages of the control (DMSO, 100%). Means ± SDs (n=3). Statistical analyses were performed using the raw data by one-way ANOVA with Bonferroni posttests. n.s., no significance; *, p< 0.05; **, p< 0.01; ***, p < 0.001. All experiments were independently repeated twice, and one representative result is presented.

Further, we investigated the effects of panobinostat, dacinostat and EBS, which are the most potent inhibitors for HPU (Figure 4A). The results showed that EBS, but not panobinostat, dacinostat or its analog AHA, has a substantial suppression on the growth of *H. pylori* (Figure 4D). The inability of AHA as well as its derivatives, i.e. panobinostat and dacinostat, on the growth of *H. pylori* as identified above seems to be consistent with the previous finding that AHA doesn’t inhibit the growth of *H. pylori*[37]. Interestingly, EBS and EBS analogs, as well as disulfiram, could dose-dependently suppress the growth of *H. pylori* and showed a minimum inhibitory concentration (MIC) in a range between 2 and 4 μg/ml (right panel of Figure 4D, Figure S6A and Table S2). Importantly, the inhibitory effect of this type of covalent inhibitors lasted for a long period in culture, as indicated by EBS and **1,** which could substantially inhibit HPU even after removal of the inhibitor for 6 h (Figure S6B).

### 3.4 Urease inhibitors prevent *H. pylori* infection in a gastric cell-based bacterial infection model

To evaluate the ability of these urease inhibitors to prevent *H. pylori* infection, we constructed a gastric cell-based bacterial infection model using the remaining viable cell number of SGC-7901 adenocarcinoma gastric cells to reflect the virulence of *H. pylori*[16]. Our results showed that treatment with 30 μM panobinostat, 30 μM dacinostat, 20 μM EBS or 20 μM disulfiram could prevent the cell death triggered by *H. pylori* (Figures 5A-B). In sharp contrast, the cells that lacked such treatments were largely sabotaged. Panobinostat and EBS were found to be the most potent agents and almost completely protected from the infection of *H. pylori.* These effects of these drugs seemed to be much more efficient than the effects of 20 μM AHA or 50 μM tinidazole, the analog of metronidazole, and one of the two antibiotics in the triple regimens for the treatment of *H. pylori* [17, 18]. In support of this observation, tinidazole as well as metronidazole hardly suppressed the growth of our *H. pylori* strain, with an MIC value of more than 512 μg/ml in culture (Figure S7A and Table S2), indicating that this strain is resistant to treatment with nitroimidazole-type antibiotics.

Since panobinostat, dacinostat, EBS and disulfiram at a concentration up to 100 μM or 25 μM did not interfere with the proliferation of SGC-7901 gastric cells (Figure S7B), the protective effects in the gastric-cell-based *H. pylori* infection model seemed to be attributed to on-targeting inhibition of the infection transmitted by *H. pylori*. Moreover, all four drugs potentially inhibited the level of ammonia in the cell medium (Figure 5C), indicating that they efficiently suppressed the endogenous urease activity of *H. pylori* in the infection model.

### 3.5 The structural basis and inhibitory mechanisms of newly-identified three classes urease inhibitors

To identify the active chemical moiety of panobinostat, dacinostat, EBS or captan required for inhibition of urease, we analyzed their structure-activity relationships (Figures 6, Table S1 and S3). The former two inhibitors are hydroxamic acid-based urease inhibitors, and not only their hydroxyamino heads are forming hydrogen bonds with the catalytic residues in JBU or HPU (Asp633 or Ala636 for JBU; Asp362 or Ala365 for HPU), but also the acetyl group constitutes one hydrogen bond (His492 for JBU and His221 for HPU; Figures 2E and S8A). Consistent with this observation, the hydroxyamino and acetyl groups of AHA interact with Asp362 or Ala365 and His221 in a co-crystal structure of AHA and HPU[2], respectively (Figure S8A). Compound lacking of this acetyl group, i.e. hydroxylamine, totally abolished the inhibitory effect of this type inhibitor (Figure 6B and Table S3). Apart from these interactions, the hydrophobic benzene ring and secondary amine group of panobinostat were found to be additional pharmacophores (upper panel, Figure 2E), which interact favorably with His492 (JBU) or His221 (HPU) and form an extra hydrogen bond with Asp494 (JBU) or Asp223 (HPU). In supporting this finding, the hydroxamic acid analogs that are lack of the benzene ring, i.e. ricolinostat, ilomastat and pracinostat, are inactive to JBU and HPU (Figure 6B and Table S3). Strikingly, the replacement of benzene with benzimidazole (pracinostat) totally loses the inhibition, suggesting the benzene is critical for maintaining the inhibition. Moreover, the secondary amine group seems to be also important for enhancing the potency of this type inhibitor, since the modification or replacement of it with hydroxyl group or sulfonyl group (dacinostat or belinostat), also weaken ∼ 5-fold or 24-fold in IC_50_ values.

**Figure 6.**
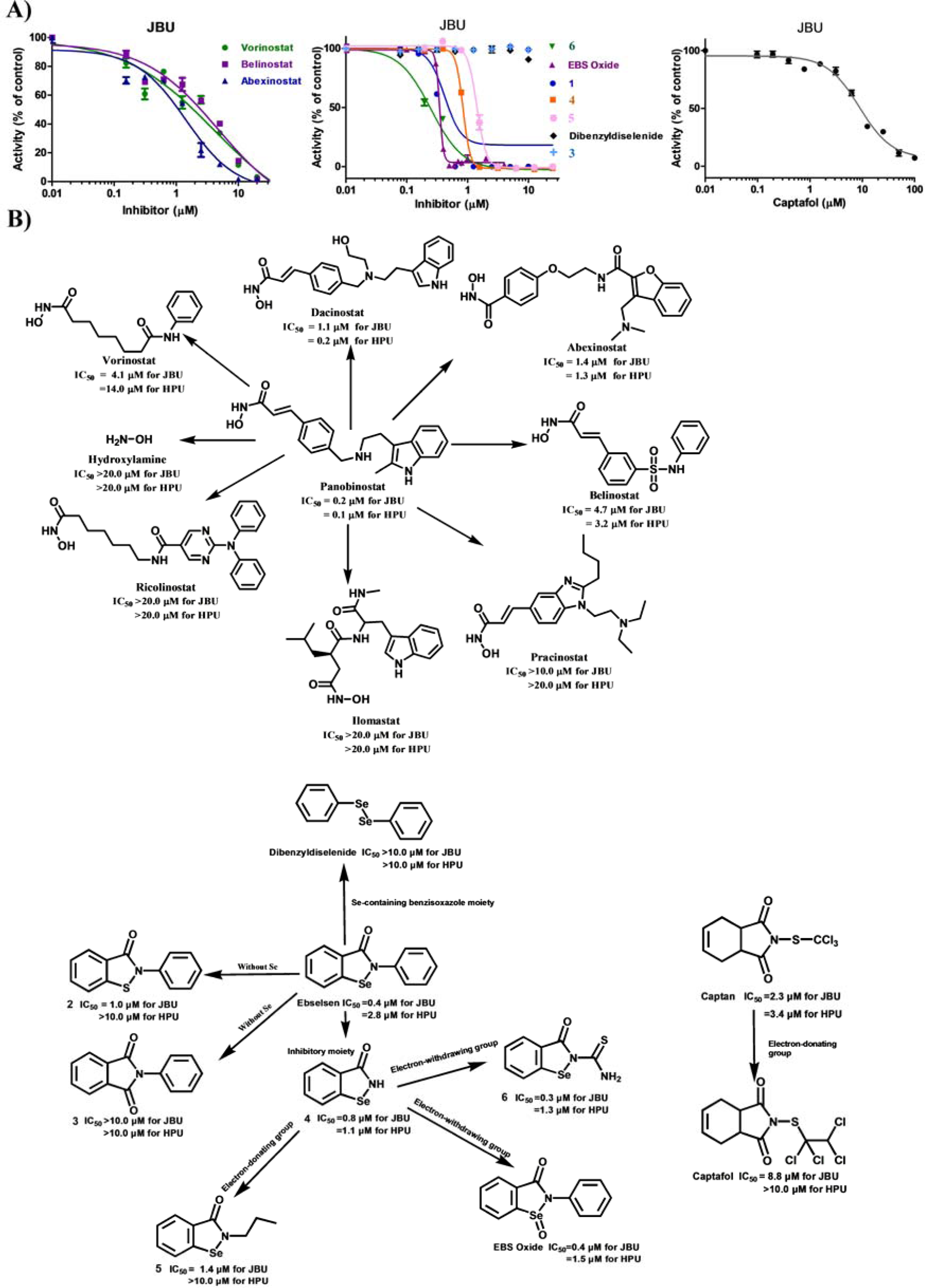
Structure-activity relationships of panobinostat, dacinostat, EBS and captan. (**A**) The effects of commercially available analogs of panobinostat and dacinostat, newly synthesized EBS derivatives and commercially available EBS or captan analogs on the activity of JBU. DMSO, 100%. Mean ± SD (n=3). The experiments were independently repeated at least twice, and one representative result is presented. (**B**) The illustration charts for the structure-activity relationships of hydroxamic acid analogs, EBS or captan.

For EBS analogs, compounds (**2**-**3**) lacking the Se atom largely lost inhibitory activities toward JBU and HPU (Figure 6B and Table S1). Furthermore, dibenzyl diselenide was also inactive toward both ureases, indicating that the Se-containing benzisoxazole moiety rather than the solo Se atom might be essential for the inhibition. Indeed, Se-containing benzisoxazole (**4**) showed potent inhibition of HPU (IC_50_ ∼ 0.8 and 1.1 μM for JBU and HPU, respectively). The introduction of an electron-donating group to the benzisoxazole moiety apparently strongly reduced the potency (**5;** IC_50_ ∼ 1.4 μM for JBU and more than 10 μM for HPU; Figure 6B). In contrast, the provision of electron-withdrawing groups to the nitrogen or Se atom of the benzisoxazole moiety, i.e., **6** or EBS oxide, seemed to enhance the potency of JBU by a maximum of three-fold (**6**). Similarly, when weakening the electron-withdrawing effect in the substitution group of the isoindole dione core of captan, the active moiety (Figure 3C), was also found to lead to a decreased potency (Figures 6; Table S1). Taken together, these data indicate that the Se-containing benzisoxazole or the isoindole dione moiety played crucial roles in the potency of these kinds of inhibitors, the Se or N atom of which was subjected to nucleophilic attack by the thiol group of Cys and formed the Se-S or N-S bond.

## 4. Discussion and conclusion

In the present study, we could identify that four clinical-used drugs, i.e., panobinostat, dacinostat, EBS and disulfiram, two anti-cancer drugs, an anti -stroke or -bipolar drugs, and an alcohol-deterrent drug, respectively, could protect the gastric cells from the infection at submicromolar concentrations (Table 1 and Figure 5). The efficacy of these drugs substantially exceeded that of AHA, a well-known urease inhibitor and clinically used drug for bacterial infections. They seemed also to be more effective than tinidazole, a metronidazole type antibiotic in the classic triple recipe for *H. pylori* (Figures 5). Moreover, panobinostat, EBS and disulfiram have been administered to humans and do not incur severe side effects[35, 38, 39]. Additionally, these drugs did not affect the viability of mammalian cells at a concentration up to 100 μM or 25 μM (Figure S7B), suggesting that they had a rather safe profile in cells and *in vivo*. Taken together, our study armed with the newly-developed HTS assay for urease repositions four clinically used drugs as new advanced leads for the treatment of *H. pylori* infection.

The mode of action of panobinostat, dacinostat, EBS or disulfiram was found to inhibit *H. pylori* urease and reduce the production of NH_3_ in culture (Table 1; Figures S6A, 4B, 4C and 5C), which are well-known bacterial virulence factors[16]. Panobinostat and dacinostat are reversible hydroxamic acid-type inhibitors for urease, and displayed more than 250 or 130 -fold potencies than its analog AHA (Table 1). These largely improved inhibitors indeed enhanced the protective effects to the infection of *H. pylori* in the cell-based infection model (Figures 5A and 5C), demonstrating that pharmacologically targeting urease could offer an effective treatment for *H. pylori* and HPU is a validated pharmacological drug target. However, suppression of the urease activity with these potent inhibitors of HPU, could not retard the growth of *H. pylori* in culture, indicating that urease is not crucial for bacterial growths.

Moreover, EBS was found to irreversibly inhibit urease by covalently modifying an allosteric Cys residue outside of the active site (Figures 2A and 3). The newly identified covalently allosteric regulation of the activity and stability of urease by EBS and captan may explain why these inhibitors could potently and persistently inhibit urease activity and the growth of *H. pylori* even in the presence of high concentrations of urea substrate (Figure S6B), two merits that are observed for covalent allosteric drugs[40]. Indeed, when compared with the reversible inhibitor AHA, EBS displayed an ∼ 400 and 10-fold improved potency for JBU and HPU, respectively, and a long-acting inhibitory effect on the endogenous activity of urease and the growth and infection of *H. pylori* in culture (Figures 4C-D, 5B-C and S6B). Importantly, the anti-*H. pylori* MIC value of EBS and its analogs, i.e. EBS oxide, **1**, **4**, **6**, seems to be much effective or at least comparable to metronidazole or clarithromycin, which are the two antibiotics in the classic triple recipe for *H. pylori* (Table S1)[41], indicating these newly-validated chemical moieties for inhibiting the growth of *H. pylori* are promising antibiotics for developing new treatments for urease-containing pathogens. Since the urease activity is dispensable for the growth of *H. pylori* (see our discussions with the mode of action of panobinostat and dacinostat), this finding indicates the effect of EBS-type inhibitor on the growth of *H. pylori* is beyond the solo inhibition of urease activity.

In summary, we identified five clinical drugs as submicromolar inhibitors for plant or bacterial urease by performing the first HTS campaign of urease. These clinically used drugs panobinostat, dacinostat, EBS and disulfiram inhibit the virulence of *H. pylori* in a gastric-cell-based infection model. This study provides a new HTS assay, drug leads and a regulatory mechanism to develop bioactive urease inhibitors for the treatment of *H. pylori* infection, especially antibiotic-resistant strains.

## Abbreviations

AHA: acetohydroxamic acid
CBS: cystathionine beta-synthase
CSE: cystathionine γ-lyase
DTT: dithiothreitol
EBS: ebselen
FAD: Foreign Approved Drugs
FDA: U.S. Food and Drug Administration
H. pylori: Helicobacter pylori
HPU: H. pylori urease
HTS: high-throughput screening
JBU: jack bean urease
K_D_: equilibrium dissociation constant
LB: Luria-Bertani liquid medium
MICs: minimum inhibitory concentrations
O. anthropi: Ochrobactrum anthropi
OAU: Ochrobactrum anthropic urease
P. mirabilis: Proteus mirabilis
SPR: surface plasmon resonance.

## CRediT authorship contribution statement

F.L., J.Y., J.Y.X., X.Y.W. and F.W. designed the study, and analyzed the data. F.Z.L. and Y.X.Z. synthesized analogs of EBS lead. Y.Y.Z. constructed the assay and performed the high-throughput screening. H.Q.F. and L.J.L. performed the LC-MS/MS analysis. Q.L. and Z.P.X. confirmed the inhibitory activity of compounds. S.S.H performed the molecular simulation. F.L., X.Y.W. and F.W. wrote the paper. All authors reviewed the results and approved the final version of the manuscript.

## Acknowledgements

This work was supported by the National Natural Science Foundation of China (31870763, 21834005), the Natural Science Foundation of Shanghai (18ZR1419500), the Shanghai Foundation for the Development of Science and Technology (19JC1413000), and the Research Fund of Medicine and Engineering of Shanghai Jiao Tong University (YG2019QNB27). We thank David Sullivan, Jun Liu and Curtis Chong of Johns Hopkins University for providing the Johns Hopkins Clinical Compound Library. We thank Prof. S.C. Tao (Shanghai Center for Systems Biomedicine, Shanghai Jiao Tong University, Shanghai, China) for kindly providing the SGC-7901 cell line. We thank Dr. J.R. Xu (Department of Radiology, Ren Ji Hospital, School of Medicine, Shanghai Jiao Tong University, Shanghai, China) for assisting with the surface plasmon resonance assay experiment.

## Appendix A. Supporting information

Supplementary data associated with this article can be found in the online version.

## Supplementary Information

### EXPERIMENTAL PROCEDURES

#### Synthesis of EBS analogs 1-6

Compound **1**-**6** were synthesized according to literature procedure(1-3), as shown in Scheme S1. The chemical reagents and solvents are purchased from commercial sources, and used without further purification, unless stated otherwise. ^1^H NMR spectra for these compounds were recorded with Bruker 400 spectrometer. The chemical shifts of ^1^H NMR spectra were referenced to tetramethylsilane (δ 0.00 ppm).

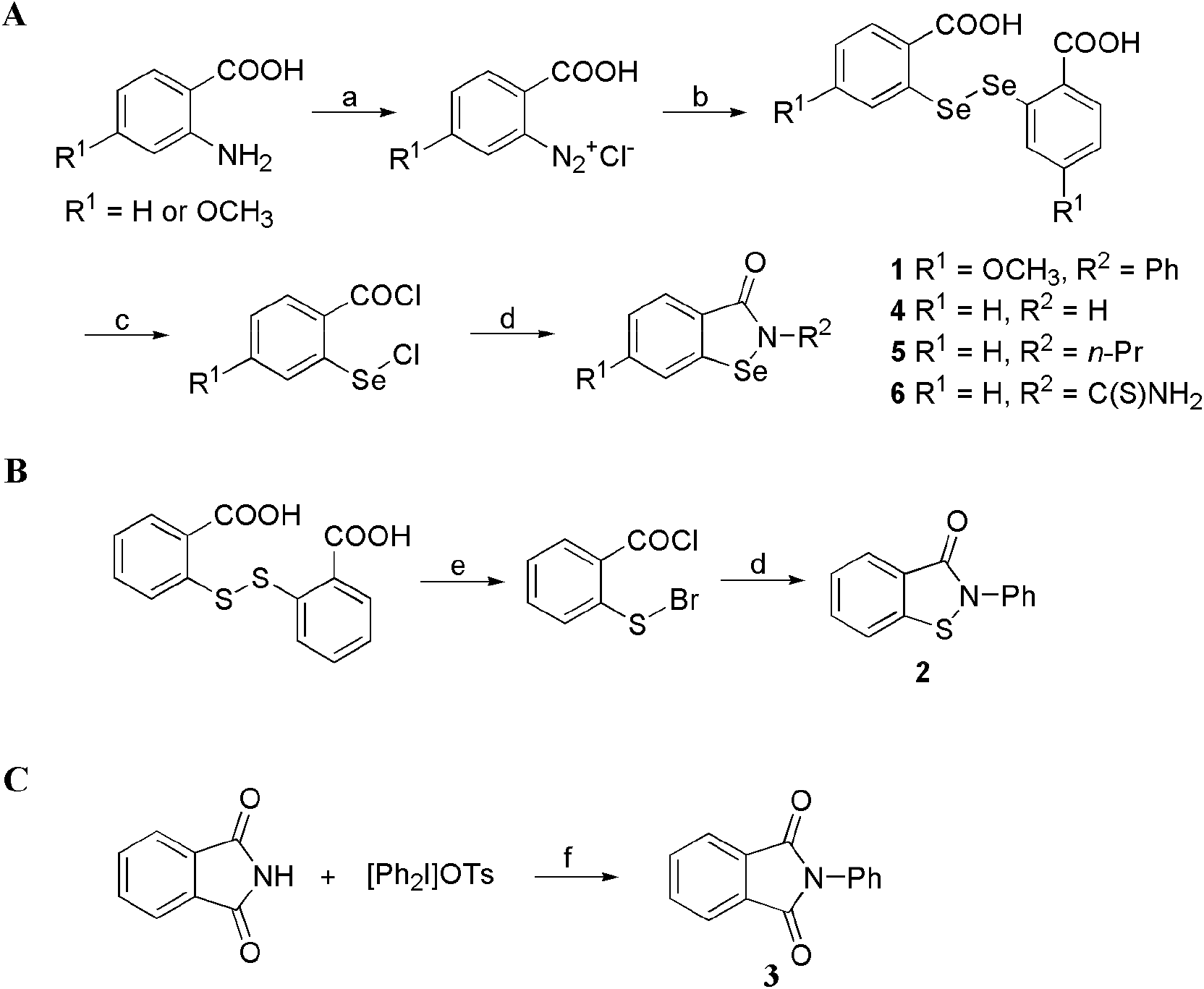

#### Scheme S1. Synthesis of compounds 1-6

Reagents and conditions: (a) HCl, NaNO_2_, 0 °C, 0.5 h; (b) Na_2_Se_2_, 60 °C, 3 h; (c) SOCl_2_, 85 °C, 3 h; (d) R^2^NH_2_, Et_3_N, CH_2_Cl_2_, rt, 4.5 h; (e) Br_2_, CH_2_Cl_2_, reflux, overnight; (f) Cu(NO_3_)_2_.xH_2_O, Et_3_N, toluene, reflux.

General procedure for synthesis of Compounds **1**, **4**, **5** and **6** (Route A).

The 2-aminobenzoic acid or its derivative was treated with hydrochloric acid (2.50 equiv.) and sodium nitrite (1.06 equiv.) in water (0.7 M) at 0 °C to form the corresponding diazonium salt. Then, the diazonium salt solution was added dropwise to a solution of Na_2_Se_2_ (0.87 equiv., fresh prepared from selenium powder and NaBH_4_ in water) at 0 °C under Argon. The stirring was continued at 60 °C for 3 h. After work-up, crude 2,2’-diseleno-dibenzoic acid was obtained. Sequentially, the acid was further converted to 2-(chloroseleno)benzoyl chloride with excess SOCl_2_ and one drop of DMF at 85 °C for 3 h. After the removal of thionyl chloride, the crude compound was obtained, and which was treated with different amines (1.2 equiv.) and Et_3_N (2.0 equiv.) in CH_2_Cl_2_ (0.1 M) under Argon to afford products **1** and **4**-**6**, respectively. Silica gel column chromatography was used to purify these compounds, and their HPLC purity was more than 99%.

2-Phenyl-6-methoxybenzoisoselen-3-one (**1**)

4-Methoxy-2-aminobenzoic acid and aniline were used to give the compound. ^1^H NMR (400 MHz, CDCl_3_): δ 8.01 (d, *J* = 8.8 Hz, 1H), 7.62 (dd, *J* = 7.6, 0.8 Hz, 2H), 7.43 (t, *J* = 8.0 Hz, 2H), 7.29-7.24 (m, 1H), 7.11 (d, *J* = 2.0 Hz, 1H), 7.01 (dd, *J* = 8.4, 2.0 Hz, 1H), 3.92 (s, 3H). MS (*m/z*): 305.0 [M+H]^+^.

Benzisoselenol-3-one (**4**)

*o*-Aminobenzoic acid and ammonia were used to give the product. ^1^H NMR (400 MHz, *d*_6_-DMSO): δ 9.17 (br, 1H), 8.06 (d, *J* = 8.1 Hz, 1H), 7.81 (dd, *J* = 8.0, 0.8 Hz, 1H), 7.61 (td, *J* = 7.6, 1.2 Hz, 1H), 7.42 (td, *J* = 7.6, 0.8 Hz, 1H). MS (*m/z*): 198.9 [M+H]^+^.

2-Propyl-benzisoselenol-3-one (**5**)

*o*-Aminobenzoic acid and *n*-propylamine were used to give the product. ^1^H NMR (400 MHz, CDCl_3_) δ 8.05 (d, *J* = 8.0 Hz, 1H), 7.63 (d, *J* = 7.6 Hz, 1H), 7.58 (td, *J* = 7.6, 1.2 Hz, 1H), 7.45-7.40 (m, 1H), 3.83 (t, *J* = 7.2 Hz, 2H), 1.76 (hex, *J* = 7.2 Hz, 2H), 1.00 (t, *J* = 7.2 Hz, 3H). MS *m/z*: 242.0 [M+H]^+^.

2-Methylthio-benzisoseleno-3-one (**6**)

*o*-Aminobenzoic acid and thiourea were used to give the product. ^1^H NMR (400 MHz, d_6_-DMSO): δ 10.21 (d, *J* = 0.8 Hz, 1H), 9.98 (d, *J* = 1.2 Hz, 1H), 8.00 (d, *J* = 8.4 Hz, 1H), 7.88 (d, *J* = 8.0 Hz, 1H), 7.71 (td, *J* = 8.0, 1.2 Hz, 1H), 7.45 (t, *J* = 7.6 Hz,1H). MS (*m/z*): 240.0 [M-NH_3_]^-^.

##### Synthesis of compound **2.**

Compound **2** was prepared according to route B (Scheme S1). 2,2’-Dithiobis-benzoic acid was reacted with bromine in CH_2_Cl_2_ under reflux and Argon, and then treated with aniline and Et_3_N in CH_2_Cl_2_ at room temperature. After purified the crude product by column chromatography, compound **2** was obtained. ^1^H NMR (400 MHz, CDCl_3_): δ 8.11 (d, *J* = 7.6 Hz, 1H), 7.73-7.69 (m, 2H), 7.68-7.65 (m, 1H), 7.51-7.43 (m, 3H), 7.59 (d, *J* = 8.0 Hz, 1H), 7.33 (t, *J* = 7.6 Hz, 1H). MS (*m/z*): 227.0 [M]^+^.

##### Synthesis of compound **3.**

Compound **3** was synthesized according to route C (Scheme S1). A Schlenk tube equipped with a stirrer bar was charged with isoindoline-1,3-dione, diphenyliododnium salt (2.05 equiv.) and Cu(NO_3_)_2_.xH_2_O (0.1 equiv.) in dry toluene (0.1 M) under Argon. The mixture was heated to 70 °C, followed by the addition of Et_3_N (1.5 equiv.). After stirring at 70 °C for 8.5 h (monitoring by TLC), the resulting mixture was continued stirring at room temperature overnight. Then, the mixture was concentrated and the residue was purified by column chromatography. ^1^H NMR (400 MHz, CDCl_3_): δ 7.99-7.94 (m, 2H), 7.80 (dd, *J* = 5.6, 3.2 Hz, 2H), 7.55-7.49 (m, 2H), 7.47-7.39 (m, 3H). MS (*m/z*): 223.1 [M]^+^.

##### HPLC method and purity analysis

The purity of compounds **1**-**5**, ebselen oxide or dibenzyl diselenide was analyzed on a Waters sunfire silica column (4.6×250mm; Waters, Milford, MA), which is coupled to a Waters HPLC system (e2695). 3 μl compound was injected onto the column and separated by a gradient elution [0 min: 95% phase A (hexane), 5% phase B (isopropyl alcohol); 15 min: 60% phase A (hexane), 40% phase B (isopropyl alcohol)] at a flow rate of 0.7 ml/min under room temperature.

Similarly, the purity of compound **6** was resolved on a Waters PHERISORB CN column (4.6×250mm, Waters). 5 μl compound **6** was injected onto the column and analyzed at a flow rate of 0.7 ml/min with an isocratic elution of solvent, which is composed of 75% hexane and 25% isopropyl alcohol.

The absorbance of the compounds were monitored at a wavelength of 230 nm, and the corresponding spectra were recorded and analyzed for the determination of the purity.

**The purity of EBS analogs, which were newly synthesized in house (Compound 1-6) or obtained from commercial sources (for Ebselen oxide and dibenzyl diselenide), were analyzed by HPLC (for details, see above).**

##### Compound 1

Determined Purity: > 99%; Retention time: 11.30 min

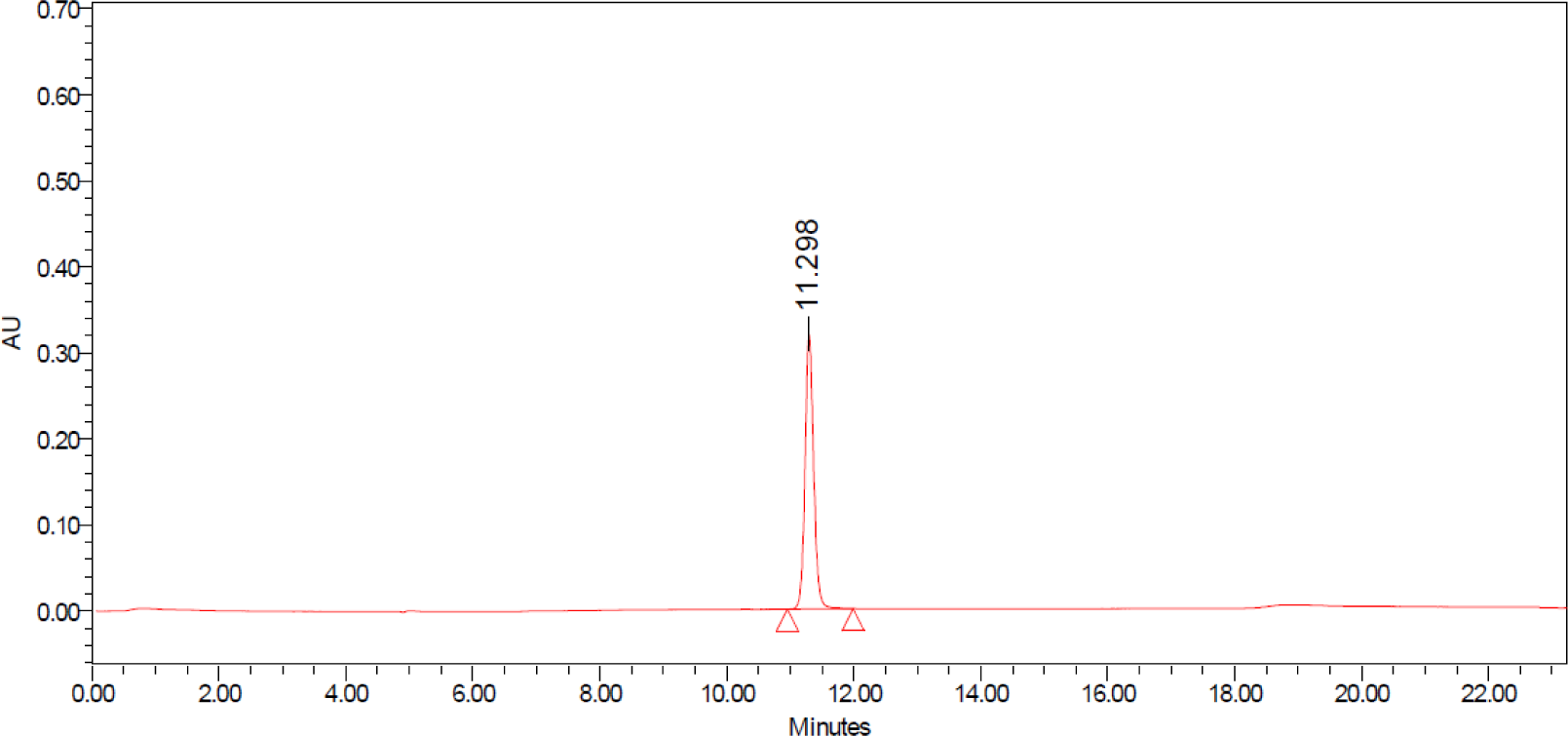

##### Compound 2

Determined Purity: > 99%; Retention time: 7.72 min

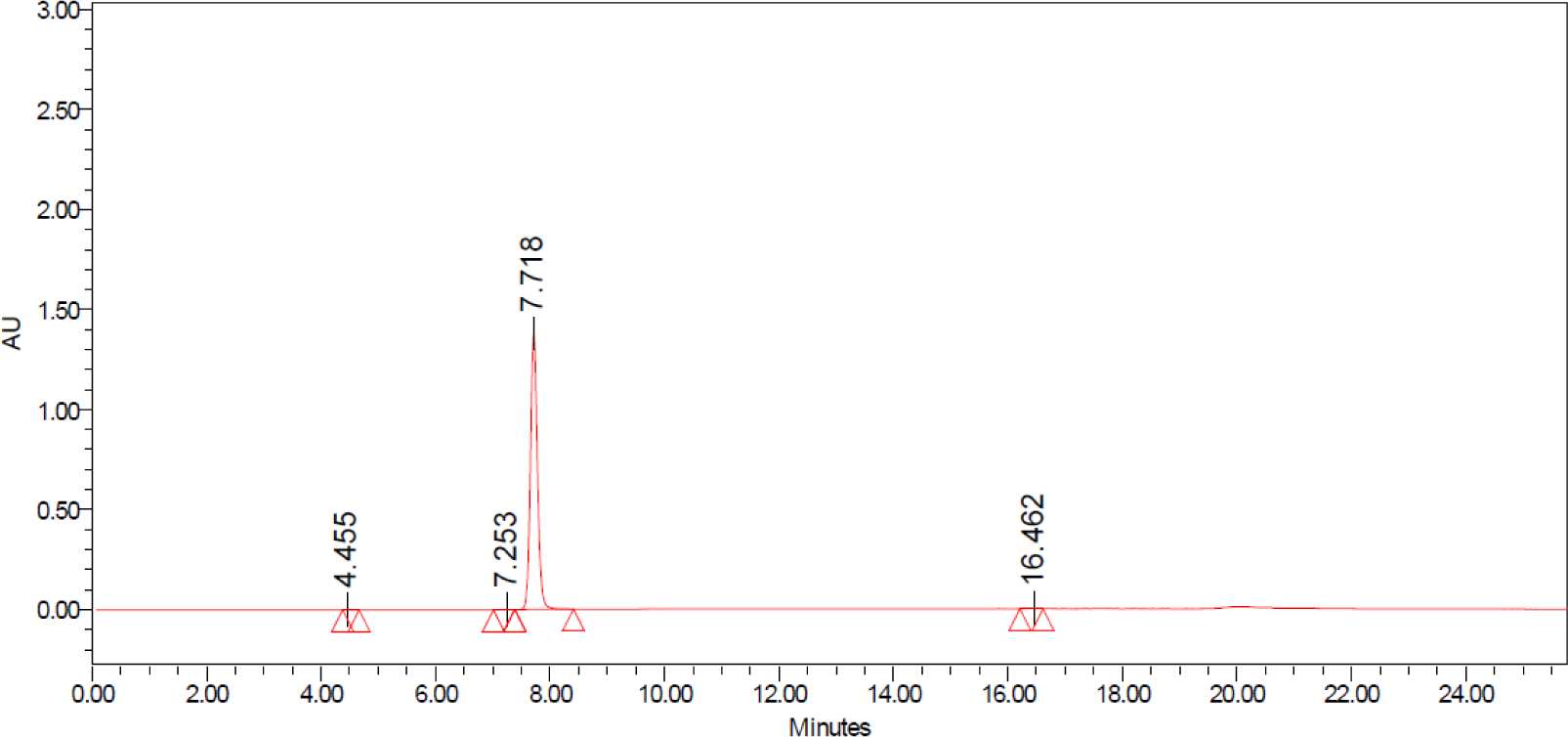

##### Compound 3

Determined Purity: > 99%; Retention time: 6.85 min

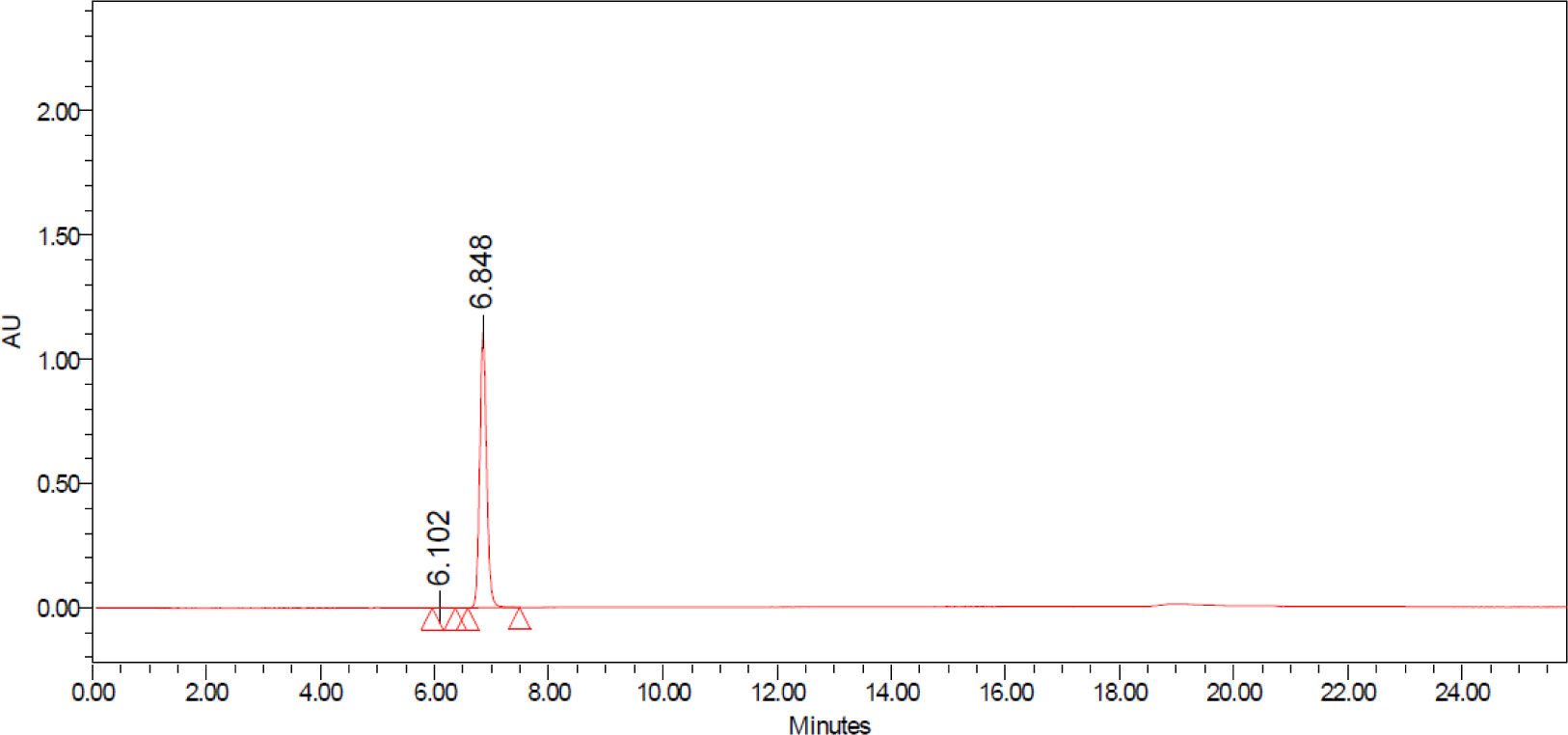

##### Compound 4

Determined Purity: > 99%; Retention time: 13.55 min

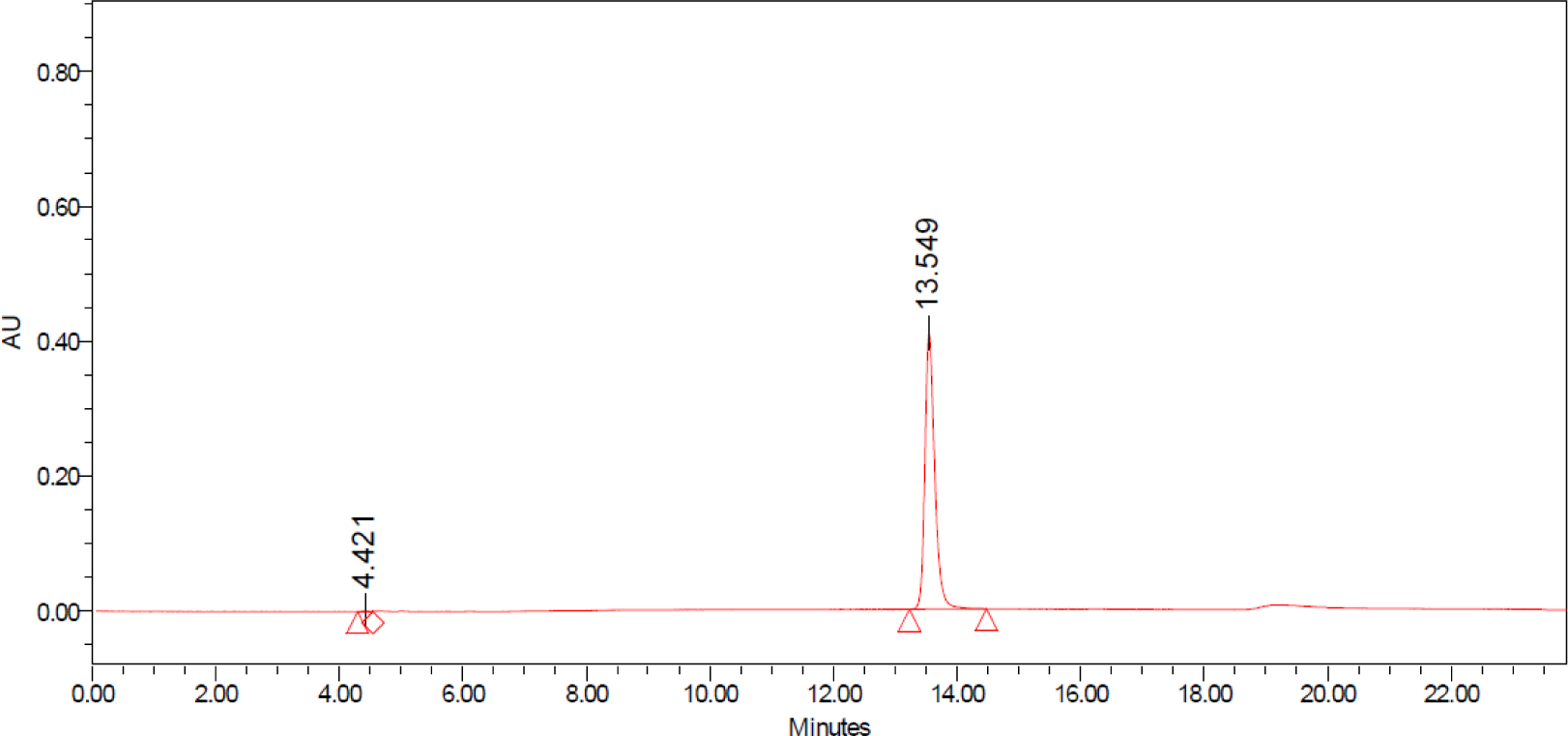

##### Compound 5

Determined Purity: > 99%; Retention time: 10.85 min

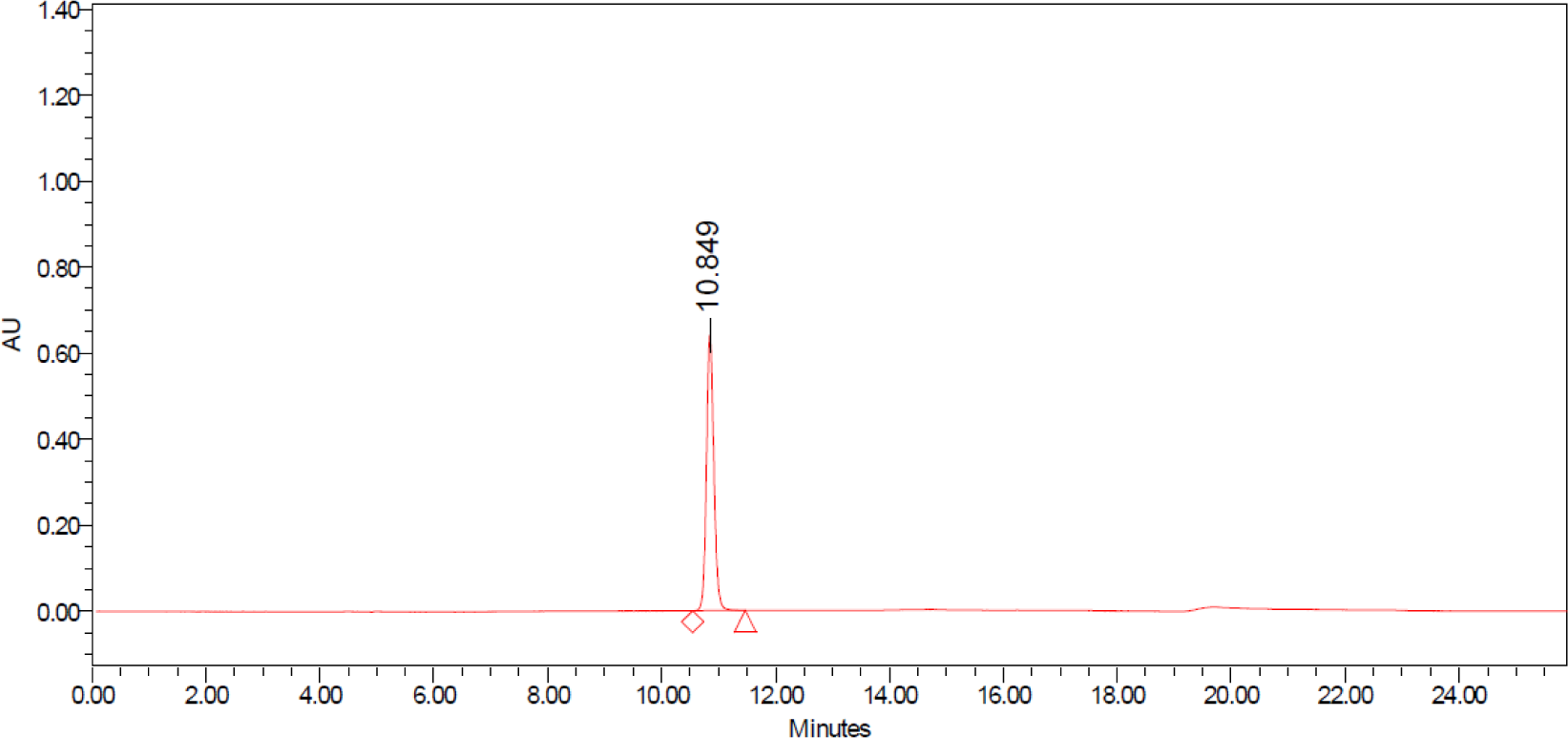

##### Compound 6

Determined Purity: > 97%; Retention time: 7.71 min

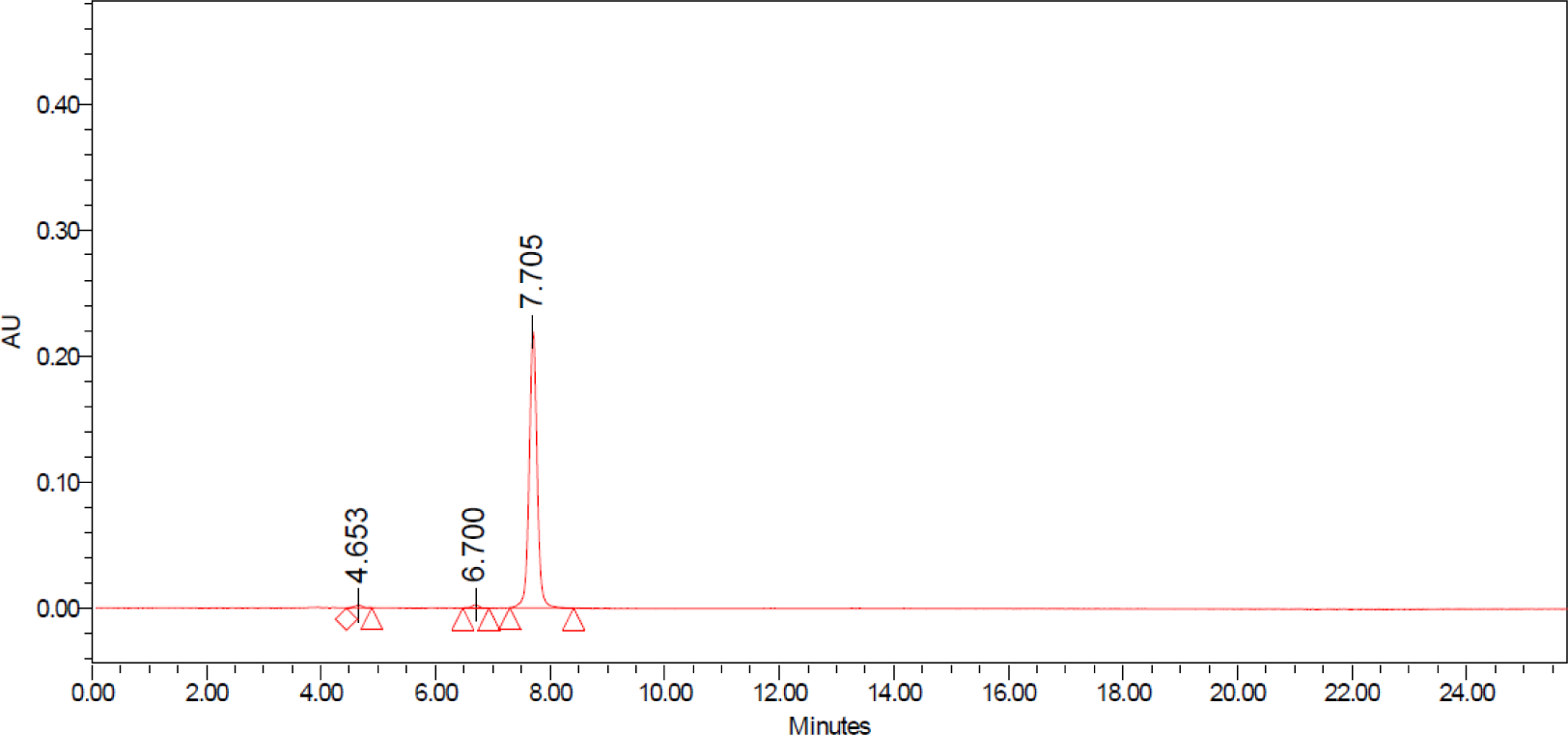

##### Ebselen oxide (Cayman)

Determined Purity: 95%; Retention time: 8.62 min

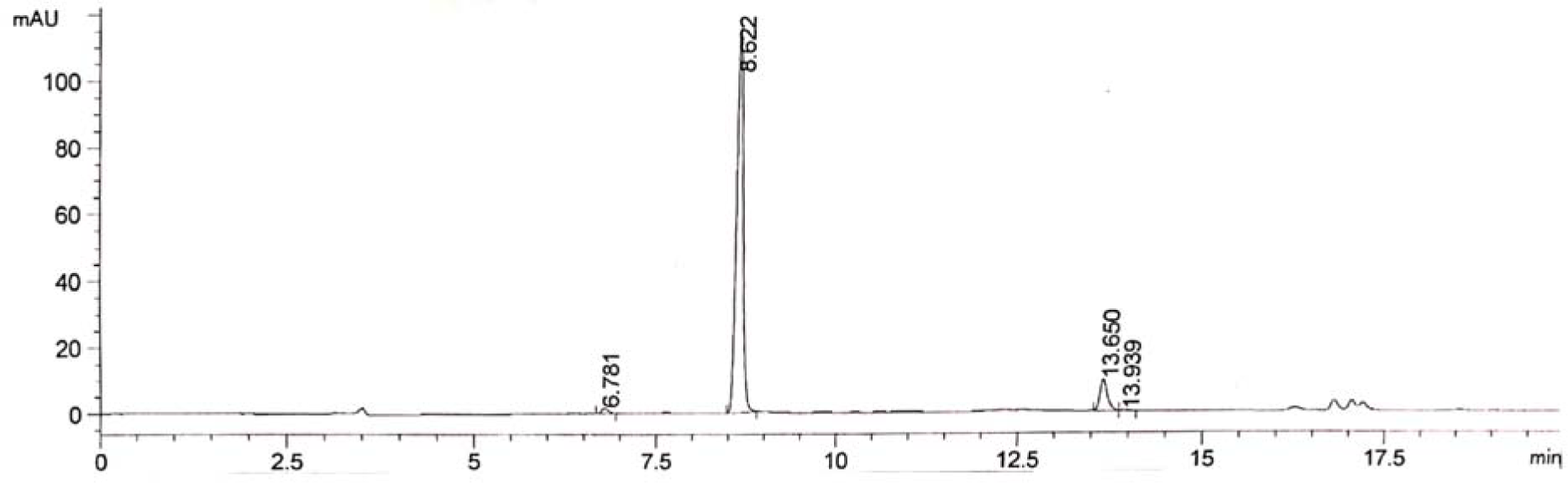

##### Ddibenzyl diselenide (Cayman)

Determined Purity: > 99%; Retention time: 4.78 min

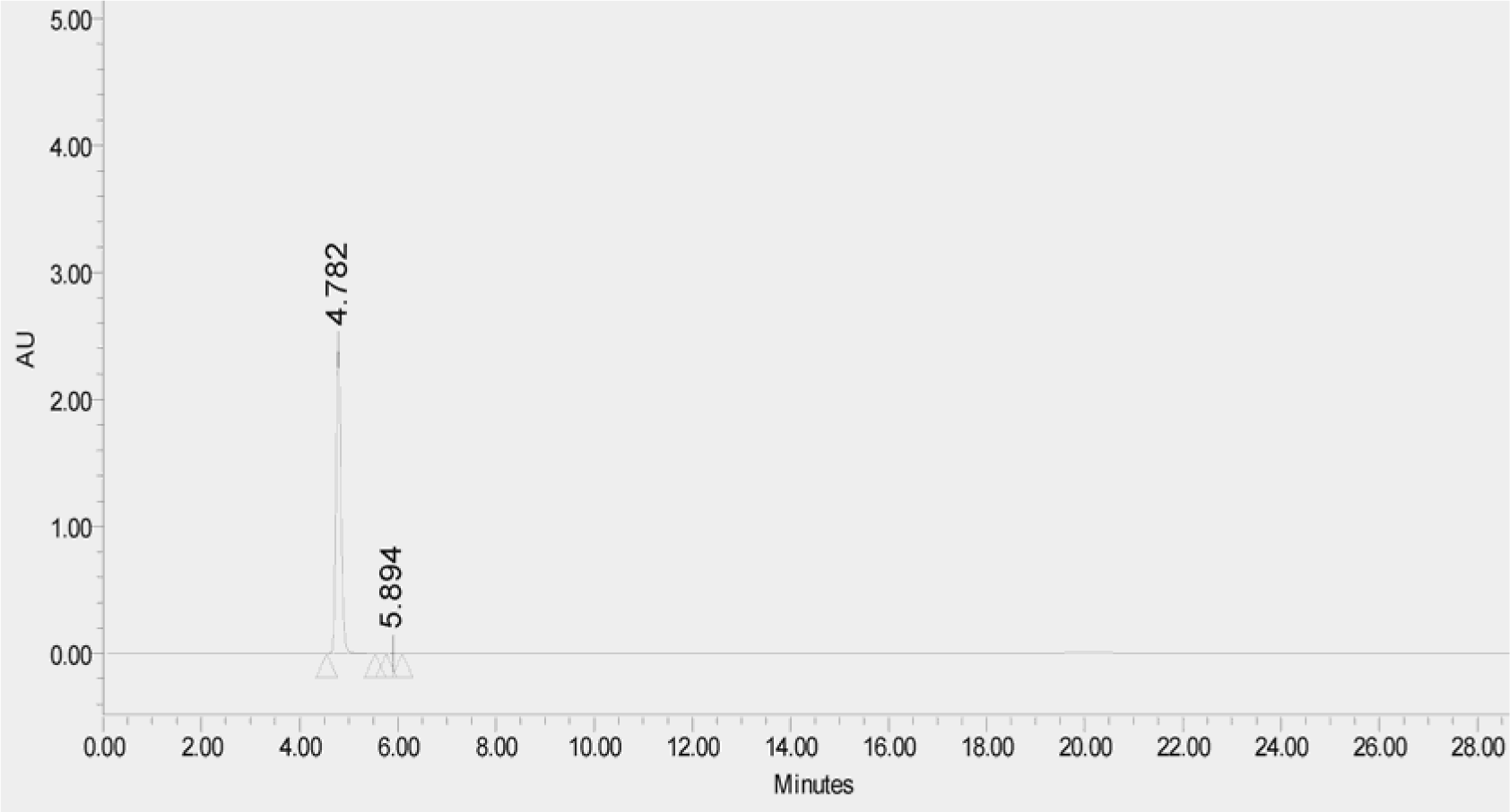

---

**Figure S1.**
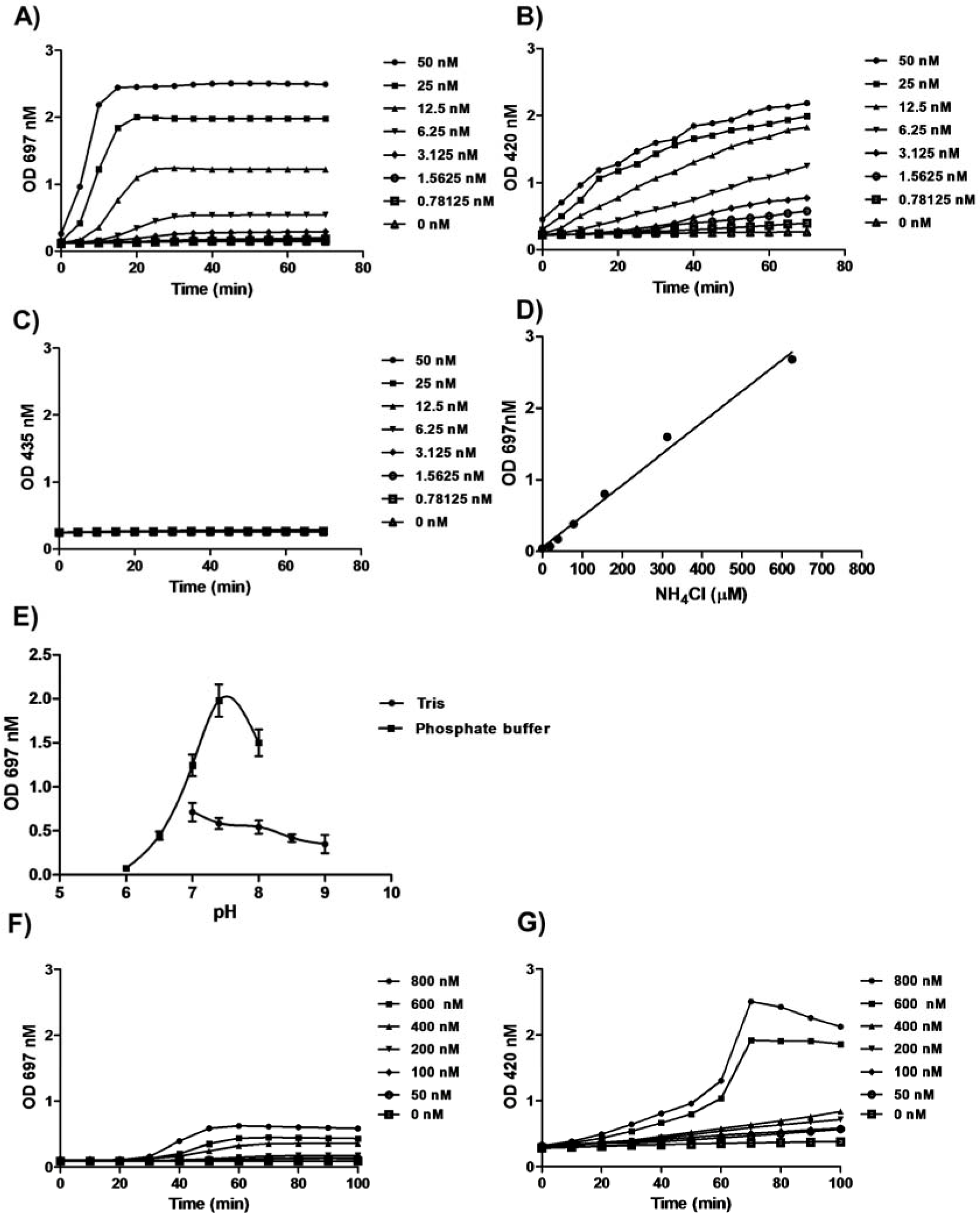
Development and optimization of the high-throughput assay for urease. Three types of detection reagents, i.e., salicylic acid-hypochlorite (**A**), Nessler’s reagent (**B**), and phenol red (**C**), were used to detect the released NH_3_ generated by JBU. The assay was monitored in the presence of various concentrations of JBU and 25 mM urea. The absorbance (O.D.) values at 697 nm, 420 nm or 435 nm were recorded accordingly. (D) Standard curve of the absorbance of indophenol blue at 697 nm versus the NH_4_Cl concentration. Various concentrations of NH_4_Cl were mixed with the detection reagent salicylic acid-hypochlorite before measurement of the absorbance at 697 nm in a microplate reader. (**E**) The pH profile of the activity of JBU. The 50 mM phosphate buffer (▪) was used to maintain the pH between 6 and 8, and 50 mM Tris-HCl (●) was used for pH 7 to 9. JBU was dissolved in the respective buffers and assayed at a final concentration of 50 nM. (**F-G**) The comparison between salicylic acid-hypochlorite and Nessler’s detection reagent for the detection of HPU activity. The assay was performed to detect the urease activity in the extract from *H. pylori* with salicylic acid-hypochlorite (left panel) and Nessler’s detection reagent (right panel) in the presence of 25 mM urea. Data are presented as the mean ± SD (n=3). The curves were fitted to the data points with GraphPad Prism 5. All the experiments were independently repeated twice, and one representative result is presented.

**Figure S2.**
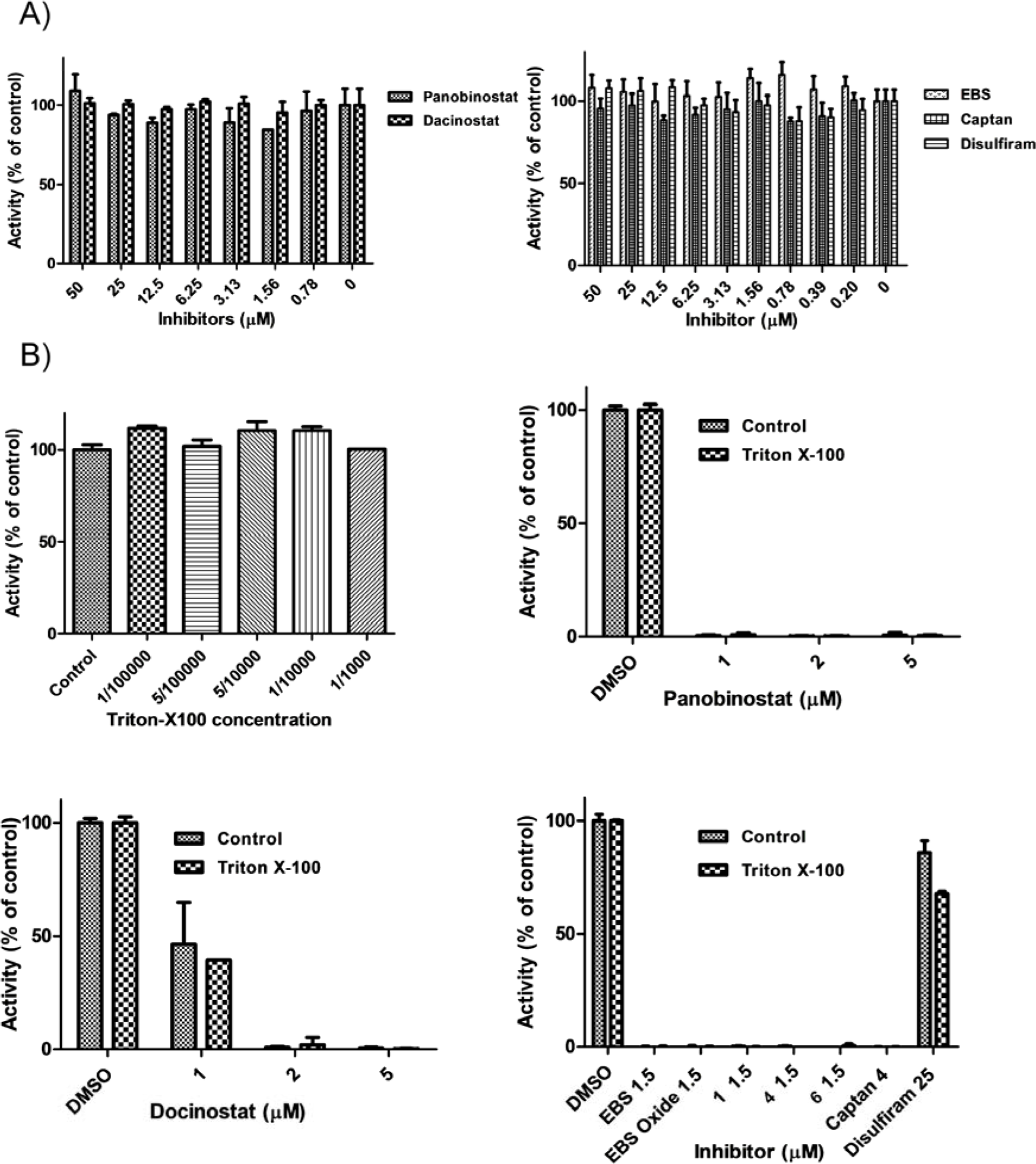
Validation of on-target inhibition of panobinostat, dacinostat, EBS, captan and disulfiram on JBU. (**A**) NH_3_ did not interfere with the inhibitors. 5 mM NH_3_·H_2_O was incubated with various concentration of panobinostat, dacinostat, EBS, captan or disulfiram in assay buffer. The volatile NH_3_ was analyzed with salicylic acid-hypochlorite detection reagent (OD_697_ nm). (**B**) Triton X-100 did not affect either the activity of JBU or the inhibition potency of panobinostat, dacinostat, EBS, captan or disulfiram as well as EBS analogs. Various concentrations of Triton X-100 were tested for their effects on the activity of JBU. Additionally, the indicated concentrations of panobinostat, dacinostat, EBS, EBS Oxide, captan, **1**, **4**, **6** or disulfiram were assayed in the presence or absence of 1/10000 Triton X-100 (v/v) to determine whether their inhibitory mechanisms occurred via colloidal aggregation (METHODS)(4). The results are shown as percentages of the respective control (DMSO or H_2_O, 100%). Mean ± SD (n=3). All experiments were independently repeated twice, and one representative result is presented.

**Figure S3.**
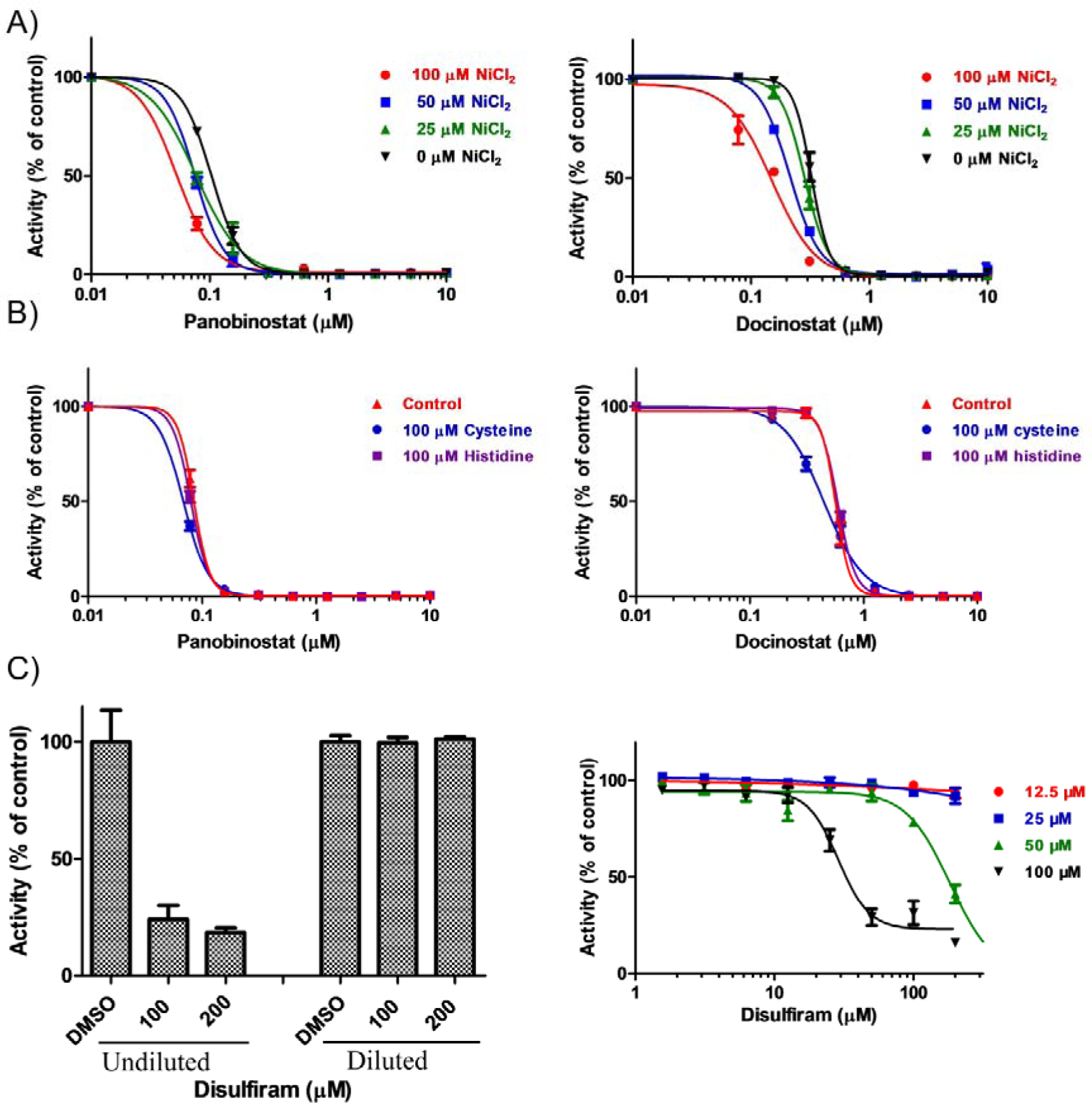
The mode of action of panobinostat, dacinostat and disulfiram *in vitro*. (**A**) The effect of NiCl_2_ on the inhibition of JBU by panobinostat or dacinostat. NiCl_2_ at a concentration of 25, 50 or 100 μM was added into the assay that is with the various concentrations of panobinostat or dacinostat under standard assay conditions. (**B**) Effects of cysteine and histidine on the inhibition of JBU with panobinostat and dacinostat. The assay samples were incubated with the indicated concentrations of panobinostat or dacinostat in the presence or the absence of 100 μM Cys or 100 μM His. The results are shown as percentages of the control (DMSO, 100%). (**C**) Reversibility of the inhibition of JBU by disulfiram. After incubation with JBU at 200, 100 μM for 60 min, disulfiram was diluted 200-fold in assay buffer. The diluted concentrations for disulfiram are 1 μM and 0.5 μM, respectively, which do not inhibit JBU (Fig. 1E). After a further incubation for 0.5 h, the remaining activity of JBU was measured accordingly (METHODS). And the effect of NiCl_2_ on the inhibition of JBU by disulfiram was shown on the right panel. The results are shown as percentages of the respective control (DMSO, 100%). Mean ± SD (n=3). All experiments were independently repeated twice, and one representative result is presented.

**Figure S4.**
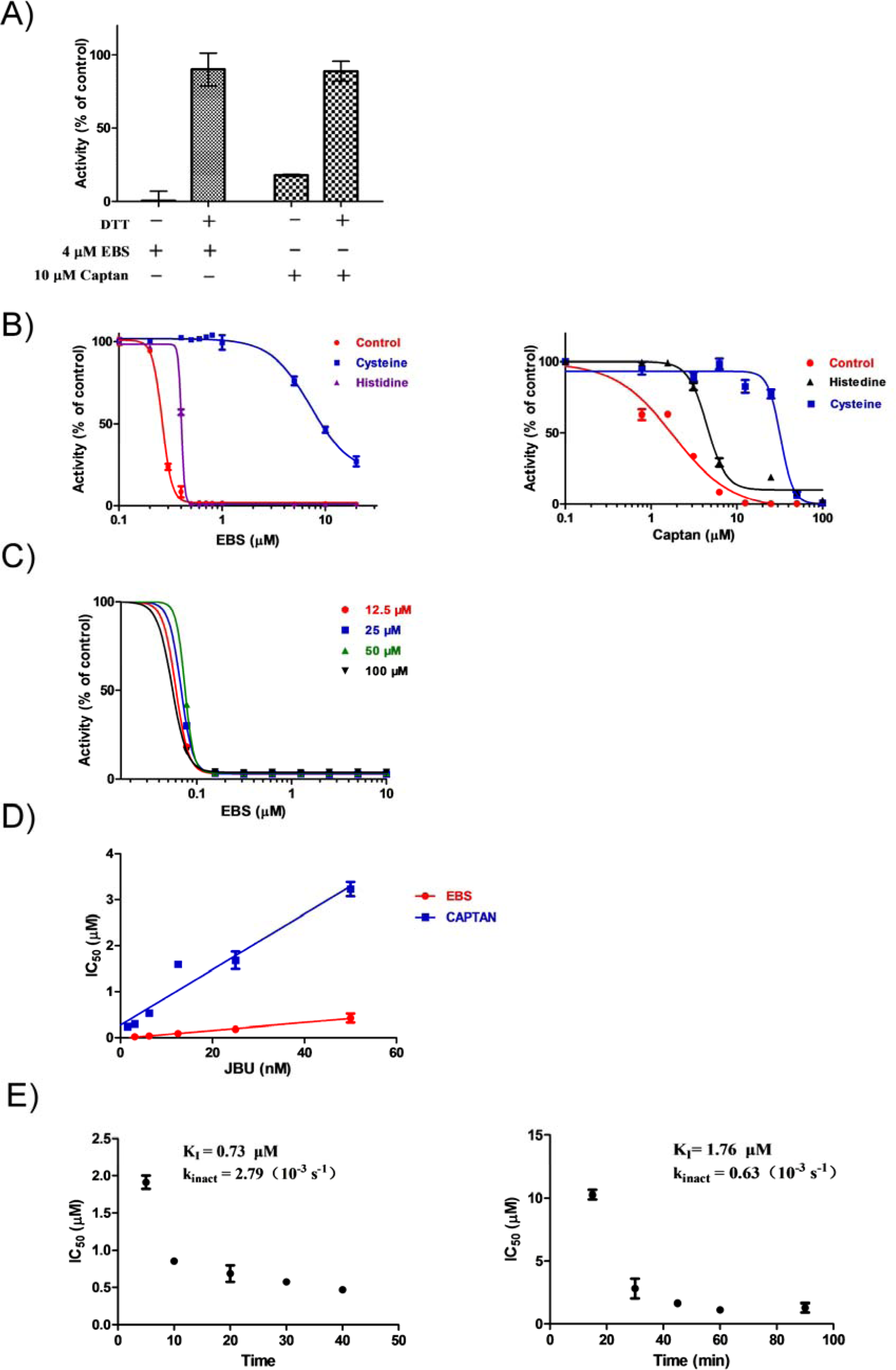
The mode of action of EBS and captan *in vitro*. (**A**) Effects of dithiothreitol on the inhibition of JBU caused by EBS and captan. The assay was incubated with 4 μM EBS or 10 μM captan in the presence or the absence of 5 mM DTT. (**B**) Effects of cysteine and histidine on the inhibition of JBU by EBS and captan. The samples were incubated with the indicated concentrations of EBS or captan in the presence or absence of 100 μM Cys or 100 μM His. (**C**) The effect of NiCl_2_ on the inhibition of EBS by JBU. NiCl_2_ at a concentration of 12.5, 25, 50 or 100 μM was incubated with the various concentrations of EBS under standard assay conditions. (**D**) The IC_50_ values of EBS and captan toward JBU were linearly correlated with the concentrations of JBU. EBS and captan were incubated with various concentrations of JBU, and the IC_50_ values were determined accordingly. (**E**) The inhibition constants of K_I_ or k_inact_ for irreversible inhibitors were determined according to the methods described in ref. (5). Means ± SDs (n=3). All experiments were independently repeated at least twice, and one representative result is presented.

**Figure S5.**
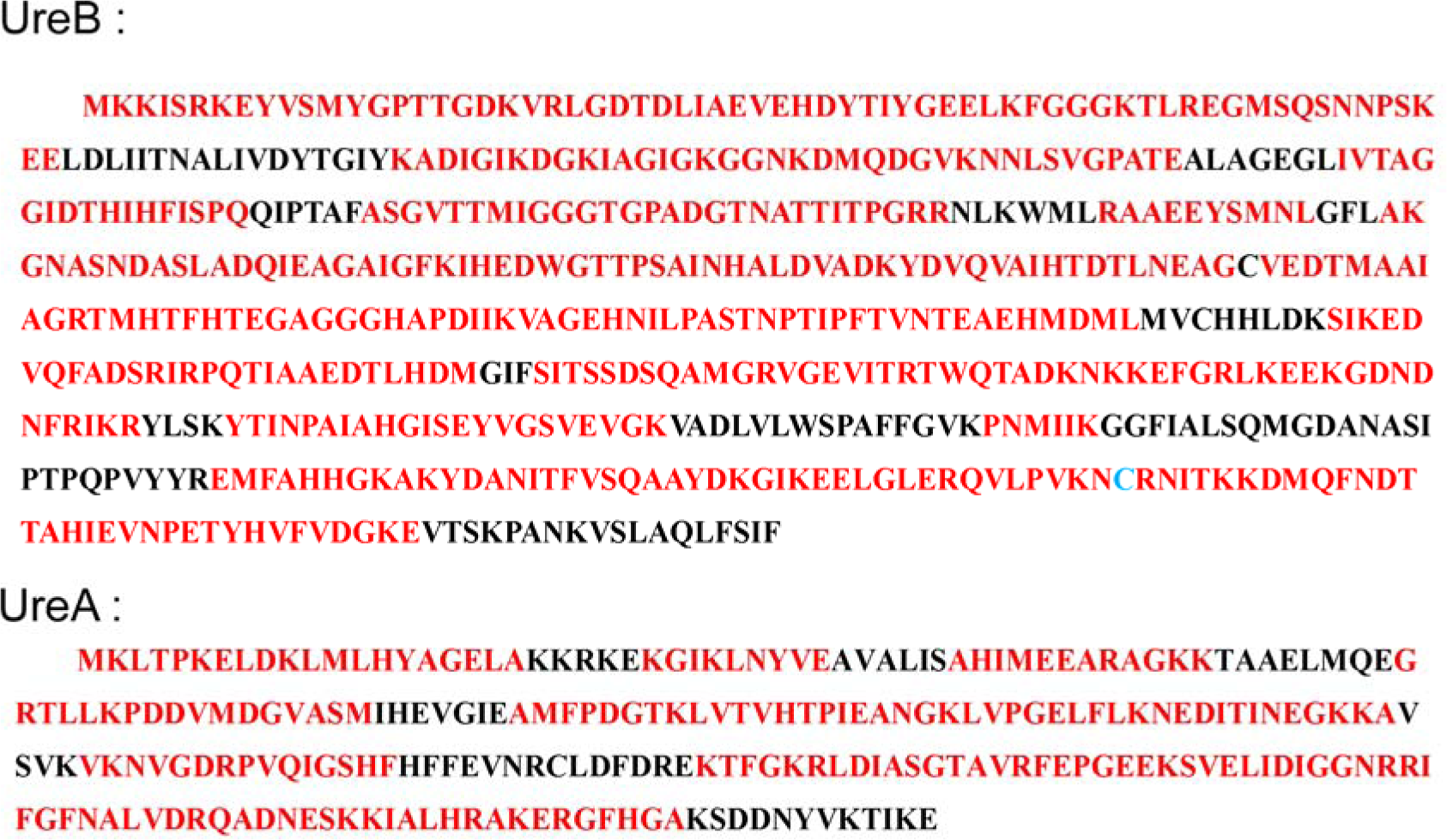
The identification of HPU from extracts of *H. pylori* by LC-MS/MS. Fraction 3 collected by size-exclusion chromatography (Figure 4B) was digested with trypsin, GluC and subtilisin, separated from the C18 reverse-phase column and subjected to analysis with a Thermo Q Exactive Orbitrap (Thermo Fisher Scientific). The peptides in red were identified by LC-MS/MS as subunit A or B of *H. pylori*. The overall coverage of UreB and UreA identified in the analysis of LC-MS/MS was 80.1% and 76.9%, respectively.

**Figure S6.**
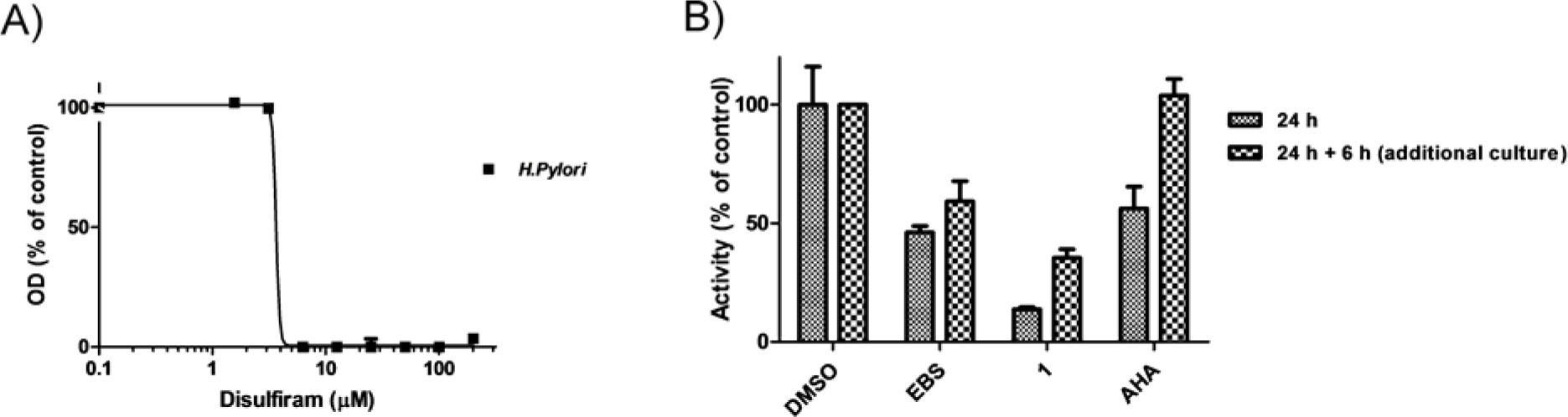
EBS and 1 is a long-acting inhibitor for HPU in culture. (**A**) Disulfiram dose-dependently and selectively inhibits the growth of *H. pylori.* Various concentrations of disulfiram were incubated at 37 °C with *H. pylori*. (**B**) The inhibitory effects of EBS and **1** on the activity of HPU *in cellulo*. EBS, **1** or AHA at a concentration of 100 μM were incubated with *H. pylori* bacteria for 24 h. Additionally, one batch of the treated bacteria was washed, diluted into freshly prepared medium without the addition of the inhibitors, and cultured for an additional 6 h. The *in cellulo* urease activities from the cultured cells under the two treated-conditions were determined accordingly (METHODS). The results are shown as percentages of the control (DMSO, 100%). Mean ± SD (n=3). All experiments were independently repeated at least twice, and one representative result is presented.

**Figure S7.**
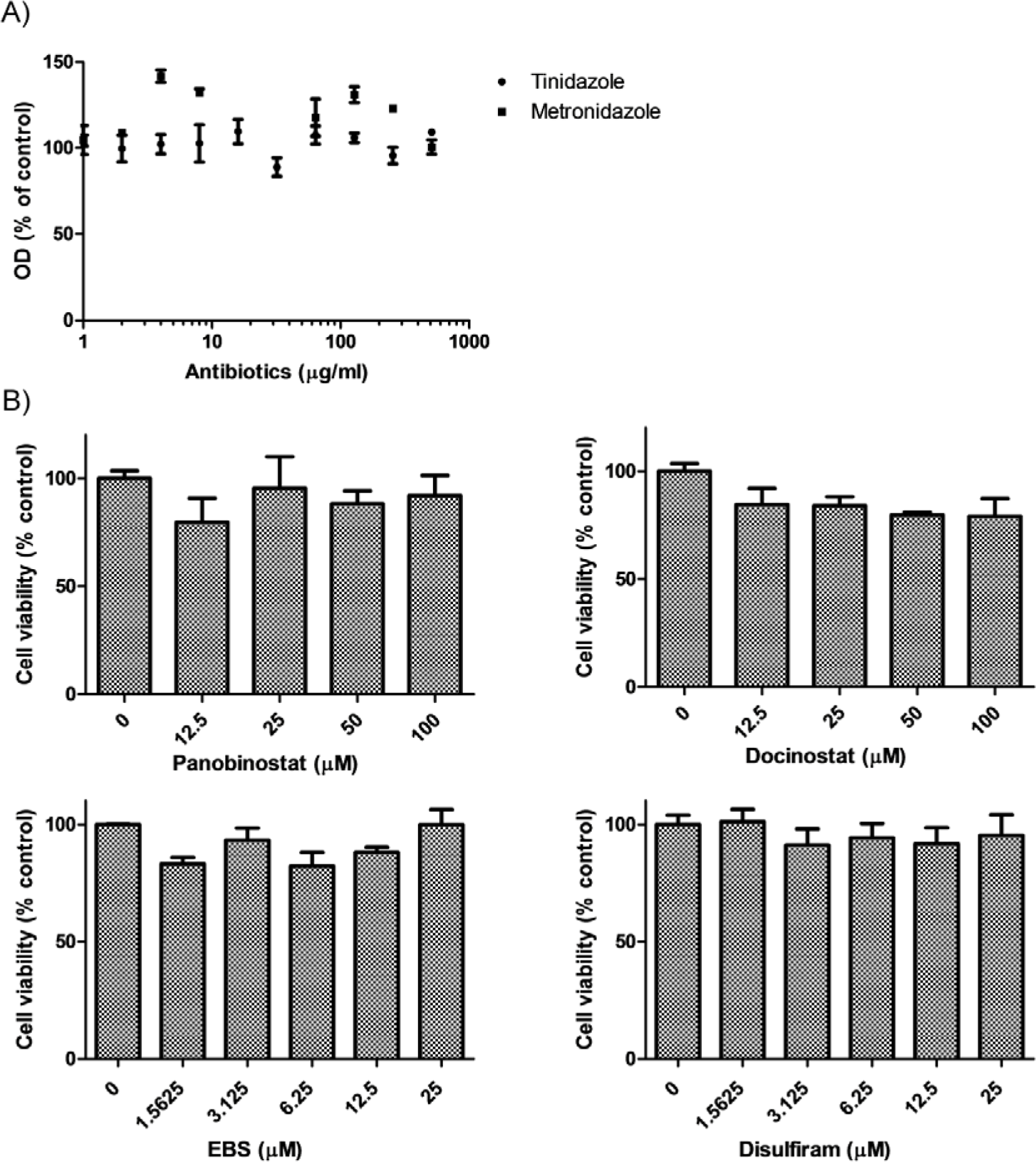
The effects of inhibitors on the cell viability of gastric SGC-7901 cells and antibiotic resistance of the *H. pylori* strain. (**A**) The *H. pylori* strain is resistant to treatment with tinidazole or metronidazole. Various concentrations of tinidazole or metronidazole were incubated at 37 °C with *H. pylori* for 72 h under standard culture conditions, and the OD at 600 nm was recorded using a spectrophotometer to determine the cell growth of *H. pylori* (METHODS). (**B**) The effects of urease inhibitors on the viability of mammalian cells. SGC-7901 cells were incubated with DMSO, the indicated concentrations of panobinosta, dacinostat, EBS or disulfiram for 24 h in a 96-well plate before measurement of cell viability using the CellTiter96® Aqueous One Solution Cell Proliferation Assay (Promega, Madison, WI). The results are shown as percentages of the control (DMSO, 100%). Means ± SDs (n=3). All experiments were independently repeated at least twice, and one representative result is presented.

**Figure S8.**
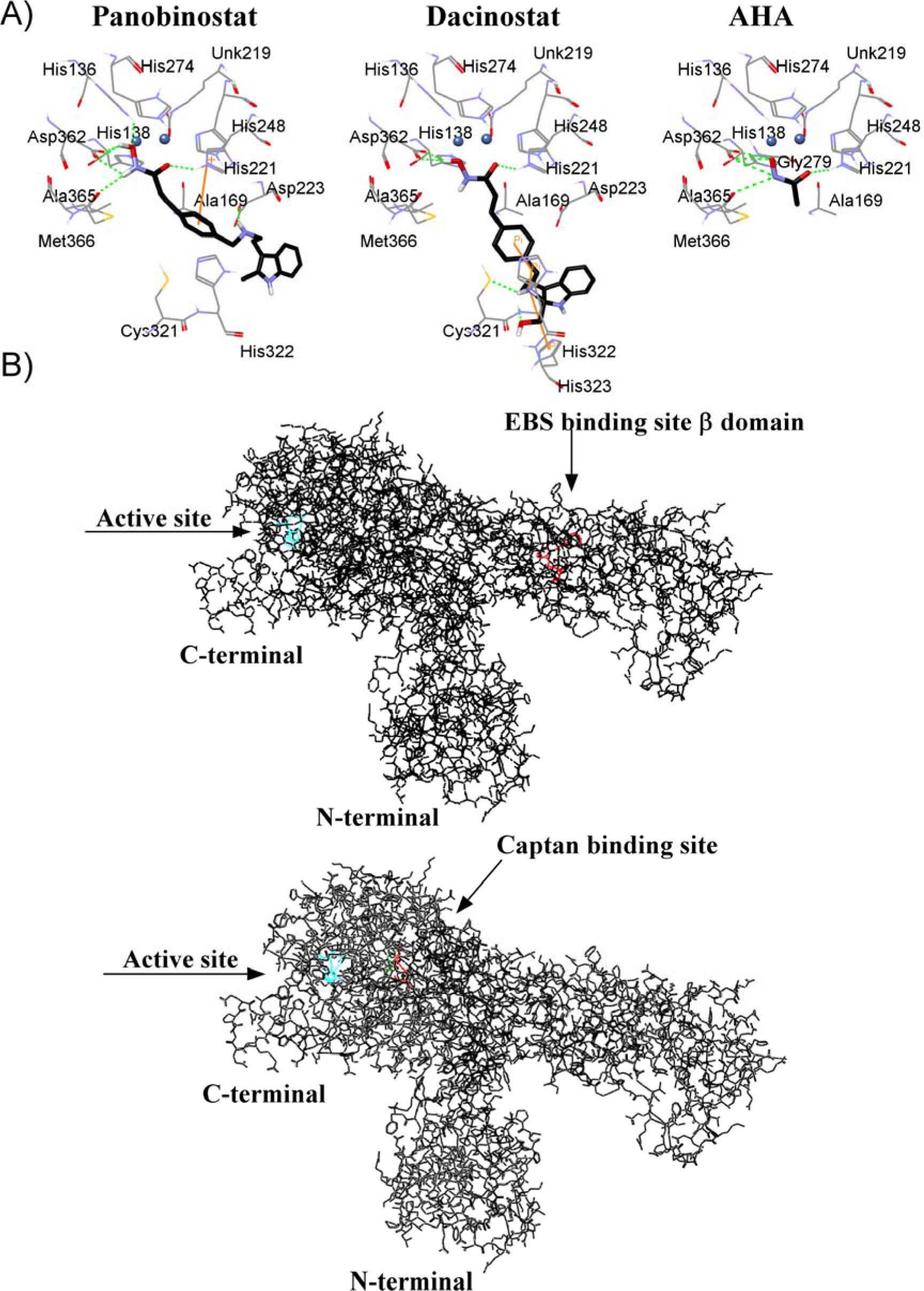
The binding modes of inhibitors in ureases. (**A**) The putative binding mode of panobinostat (black) or dacinostat (black) in the HPU active site. Panobinostat and dacinostat were docked into the HPU crystal structure (PDB code 1E9Y; ref. (6)) using the Discovery Studio software. Residues surrounding the inhibitor within a distance of 3.5 Å are shown in gray or in the default atom color. (**B**) Global view of the binding region of EBS (upper panel) and captan (lower) in JBU. In the modeled EBS or captan and protein complex structure (METHODS and Figure 3E), the protein is shown in black, the key residues (His492 and His519) in the active site of JBU in cyan and the inhibitors as well as its attached Cys residue (Cys313 for EBS, Cys406 for captan; Figure 3E) in red. Hydrogen bonds are represented as green dotted lines.

**Table S1.**
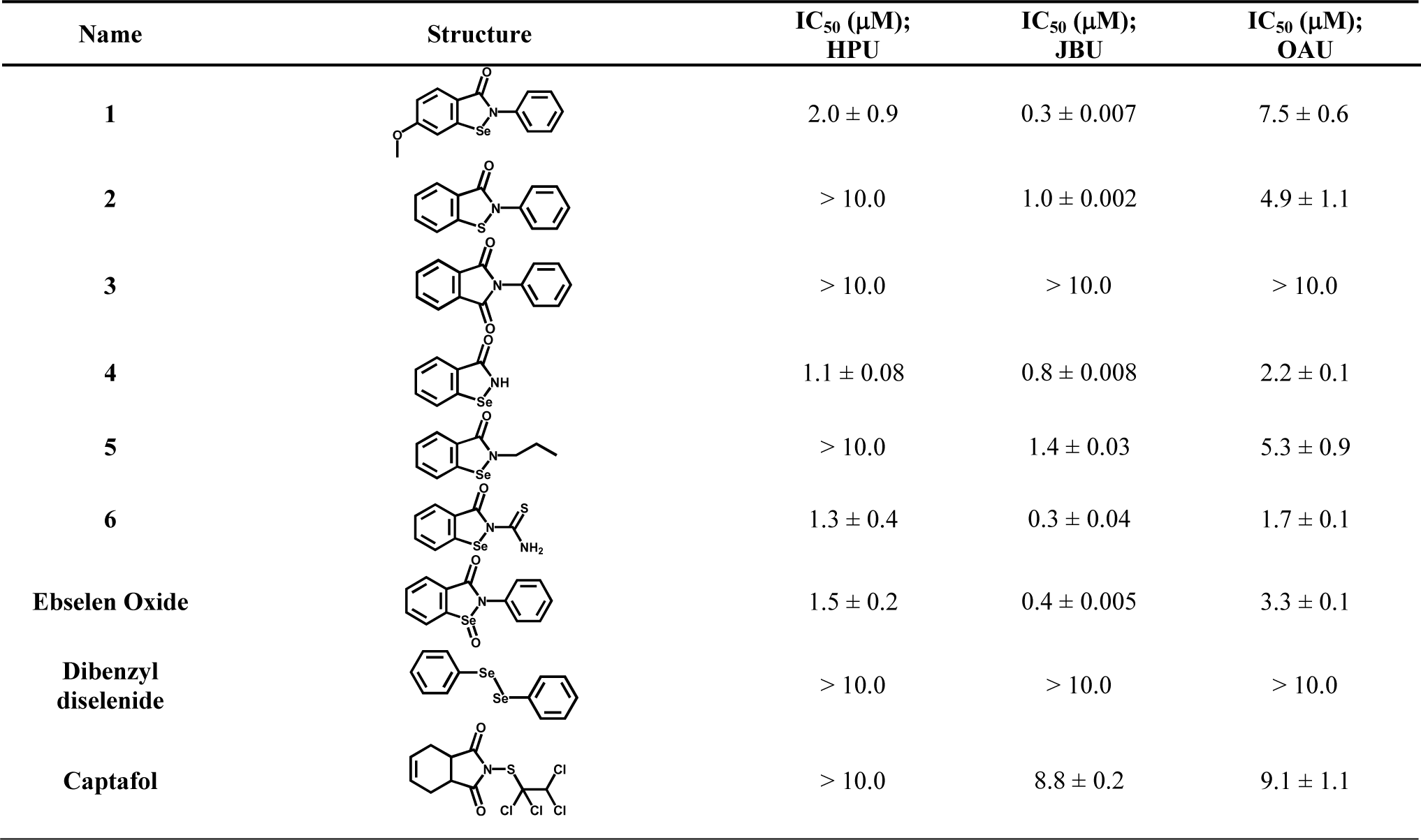
Chemical structures and IC_50_ values of EBS or captan analogs for ureases.

| Name | Structure | IC<sub>50</sub> (μM);<br>HPU | IC<sub>50</sub> (μM);<br>JBU | IC<sub>50</sub> (μM);<br>OAU |
| --- | --- | --- | --- | --- |
| 1 |  | 2.0 ± 0.9 | 0.3 ± 0.007 | 7.5 ± 0.6 |
| 2 |  | > 10.0 | 1.0 ± 0.002 | 4.9 ± 1.1 |
| 3 |  | > 10.0 | > 10.0 | > 10.0 |
| 4 |  | 1.1 ± 0.08 | 0.8 ± 0.008 | 2.2 ± 0.1 |
| 5 |  | > 10.0 | 1.4 ± 0.03 | 5.3 ± 0.9 |
| 6 |  | 1.3 ± 0.4 | 0.3 ± 0.04 | 1.7 ± 0.1 |
| Ebselen Oxide |  | 1.5 ± 0.2 | 0.4 ± 0.005 | 3.3 ± 0.1 |
| Dibenzyl diselenide |  | > 10.0 | > 10.0 | > 10.0 |
| Captafol |  | > 10.0 | 8.8 ± 0.2 | 9.1 ± 1.1 |

**Table S2.** The minimal inhibitory concentration of urease inhibitors or known antibiotics for inhibiting *H. pylori* and their IC_50_ values in the i*n cellulo* urease assay.

| Compound | *H. pylori* (MIC) | | *H. pylori* (IC<sub>50</sub> values in the *in cellulo* urease assay; μM) |
| --- | --- | --- | --- |
| | μg/ml | μM | |
| **EBS** | 4 | 12.5 | 5.7 ± 1.3 |
| **1** | 2 | 6.25 | 4.7 ± 1.1 |
| **4** | 2 | 12.5 | 18.5 ± 1.2 |
| **6** | 4 | 12.5 | 21.8 ± 1.0 |
| **EBS Oxide** | 4 | 12.5 | 23.2 ± 1.1 |
| **Captan** | 32 | 100 | 29.5 ± 1.2 |
| **Disulfiram** | 4 | 12.5 | 36.3 ± 1.0 |
| **Dibenzyl diselenide** | > 64 | > 200 | > 200.0 |
| **AHA** | > 16 | > 100 | - |
| **Tinidazole** | > 512 | > 2000 | - |
| **Metronidazole** | > 512 | > 3000 | - |

MIC; minimal inhibitory concentration

**Table S3.**
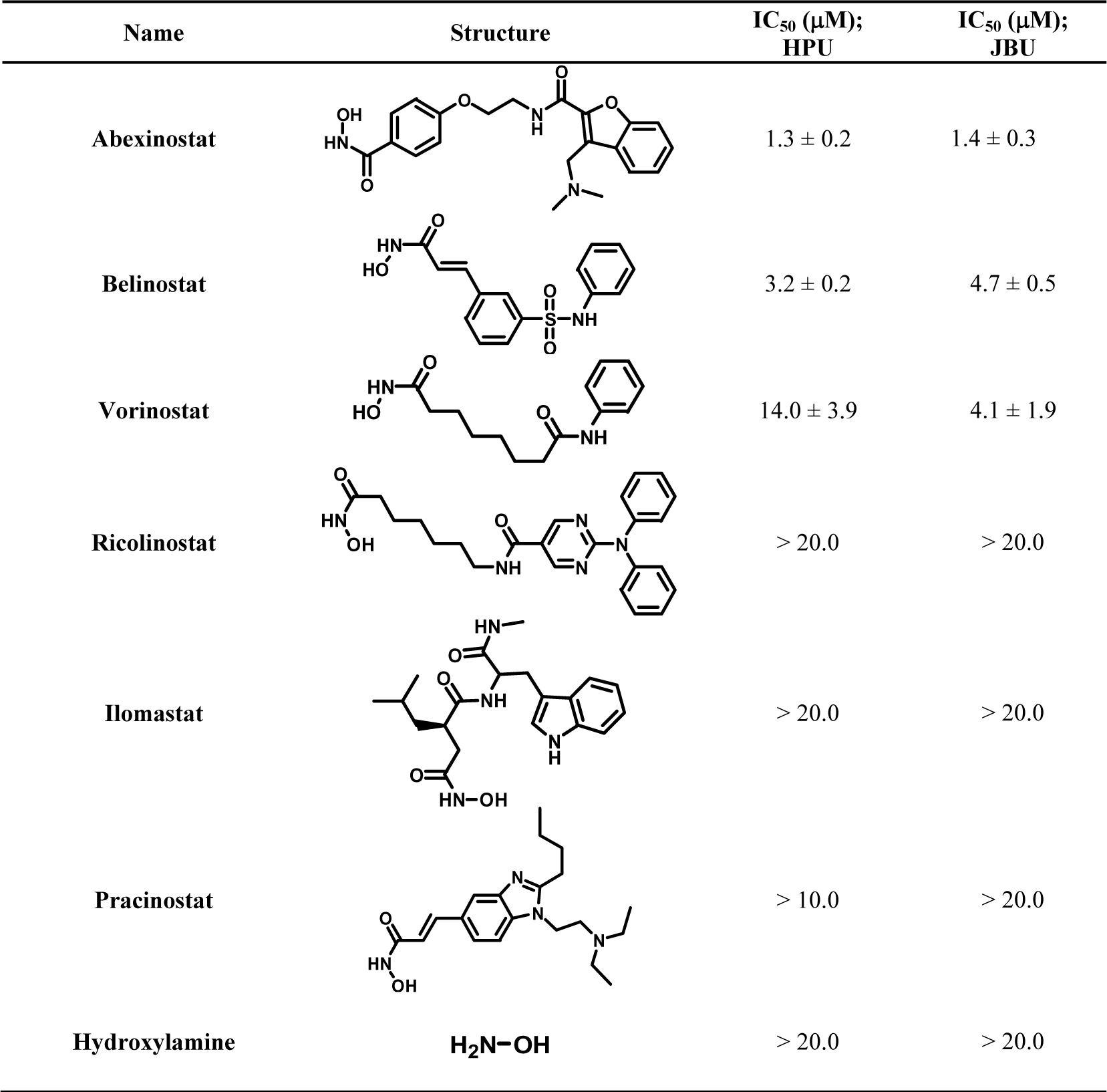
Chemical structures and IC_50_ values of hydroxamic acid-based analogs for ureases.

| Name | Structure | IC<sub>50</sub> (μM);<br>HPU | IC<sub>50</sub> (μM);<br>JBU |
| --- | --- | --- | --- |
| Abexinostat |  | 1.3 ± 0.2 | 1.4 ± 0.3 |
| Belinostat |  | 3.2 ± 0.2 | 4.7 ± 0.5 |
| Vorinostat |  | 14.0 ± 3.9 | 4.1 ± 1.9 |
| Ricolinostat |  | > 20.0 | > 20.0 |
| Ilomastat |  | > 20.0 | > 20.0 |
| Pracinostat |  | > 10.0 | > 20.0 |
| Hydroxylamine | H<sub>2</sub>N-OH | > 20.0 | > 20.0 |

**Table S4.**
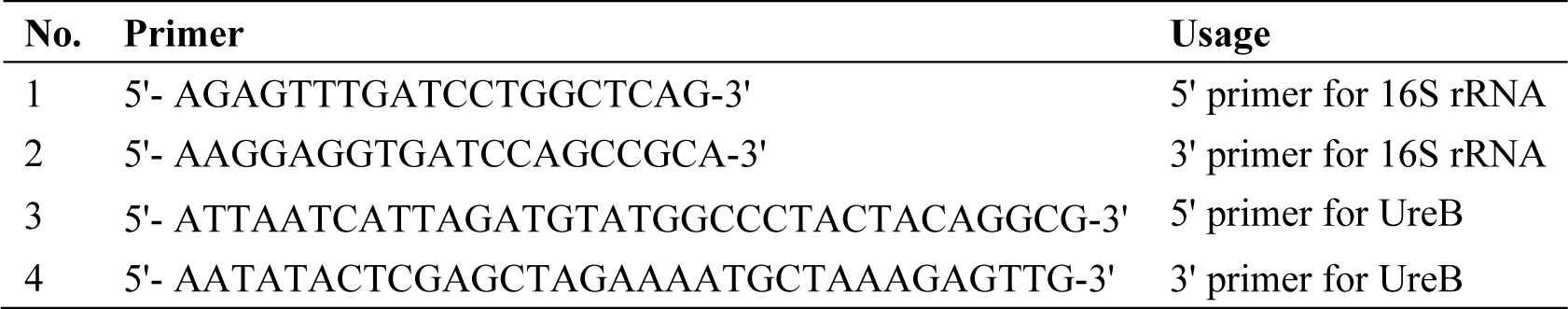
Primer sequences.

